# Integrating molecular and clinical variables to predict myocardial recovery

**DOI:** 10.1101/2024.04.16.589326

**Authors:** Joseph R. Visker, Ben J. Brintz, Christos P. Kyriakopoulos, Yanni Hillas, Iosif Taleb, Rachit Badolia, Thirupura S. Shankar, Junedh M. Amrute, Jing Ling, Rana Hamouche, Eleni Tseliou, Sutip Navankasattusas, Omar Wever-Pinzon, Gregory S. Ducker, William L. Holland, Scott A. Summers, Steven C. Koenig, Thomas C. Hanff, Kory Lavine, Srinivas Murali, Stephen Bailey, Rami Alharethi, Craig H. Selzman, Palak Shah, Mark S. Slaughter, Manreet Kanwar, Stavros G. Drakos

**Affiliations:** Nora Eccles Harrison Cardiovascular Research and Training Institute, University of Utah, Salt Lake City, UT, USA; Utah Transplantation Affiliated Hospitals (U.T.A.H.) Cardiac Transplant Program & Utah Cardiac Recovery (UCAR) Program (University of Utah Health & School of Medicine, Intermountain Medical Center, Salt Lake City VA Medical Center), Salt Lake City, UT, USA; Division of Epidemiology, University of Utah, Salt Lake City, UT, USA; Cardiovascular Division, Department of Medicine, Washington University School of Medicine, St. Louis, MO, USA; Department of Biochemistry, University of Utah, Salt Lake City, UT, USA; Department of Nutrition and Integrative Physiology and the Diabetes and Metabolism Research Center, University of Utah, Salt Lake City, UT, USA; Departments of Bioengineering and Cardiothoracic Surgery, University of Louisville, Louisville, KY, USA; Cardiovascular Institute, Allegheny Health Network, Pittsburgh, PA, USA; Heart Failure, Mechanical Circulatory Support and Transplant, Inova Heart and Vascular Institute, VA, USA

**Keywords:** Myocardial Recovery, Mechanical Circulatory Support, Heart Failure, Reverse Remodeling, Transcriptomics, Clinical Characteristics

## Abstract

Mechanical unloading and circulatory support with left ventricular assist devices (LVADs) mediate significant myocardial improvement in a subset of advanced heart failure (HF) patients. The clinical and biological phenomena associated with cardiac recovery are under intensive investigation. Left ventricular (LV) apical tissue, alongside clinical data, were collected from HF patients at the time of LVAD implantation (n=208). RNA was isolated and mRNA transcripts were identified through RNA sequencing and confirmed with RT-qPCR. To our knowledge this is the first study to combine transcriptomic and clinical data to derive predictors of myocardial recovery. We used a bioinformatic approach to integrate 59 clinical variables and 22,373 mRNA transcripts at the time of LVAD implantation for the prediction of post-LVAD myocardial recovery defined as LV ejection fraction (LVEF) ≥40% and LV end-diastolic diameter (LVEDD) ≤5.9cm, as well as functional and structural LV improvement independently by using LVEF and LVEDD as continuous variables, respectively. To substantiate the predicted variables, we used a multi-model approach with logistic and linear regressions. Combining RNA and clinical data resulted in a gradient boosted model with 80 features achieving an AUC of 0.731±0.15 for predicting myocardial recovery. Variables associated with myocardial recovery from a clinical standpoint included HF duration, pre-LVAD LVEF, LVEDD, and HF pharmacologic therapy, and *LRRN4CL* (ligand binding and programmed cell death) from a biological standpoint. Our findings could have diagnostic, prognostic, and therapeutic implications for advanced HF patients, and inform the care of the broader HF population.

**GRAPHICAL ABSTRACT:** 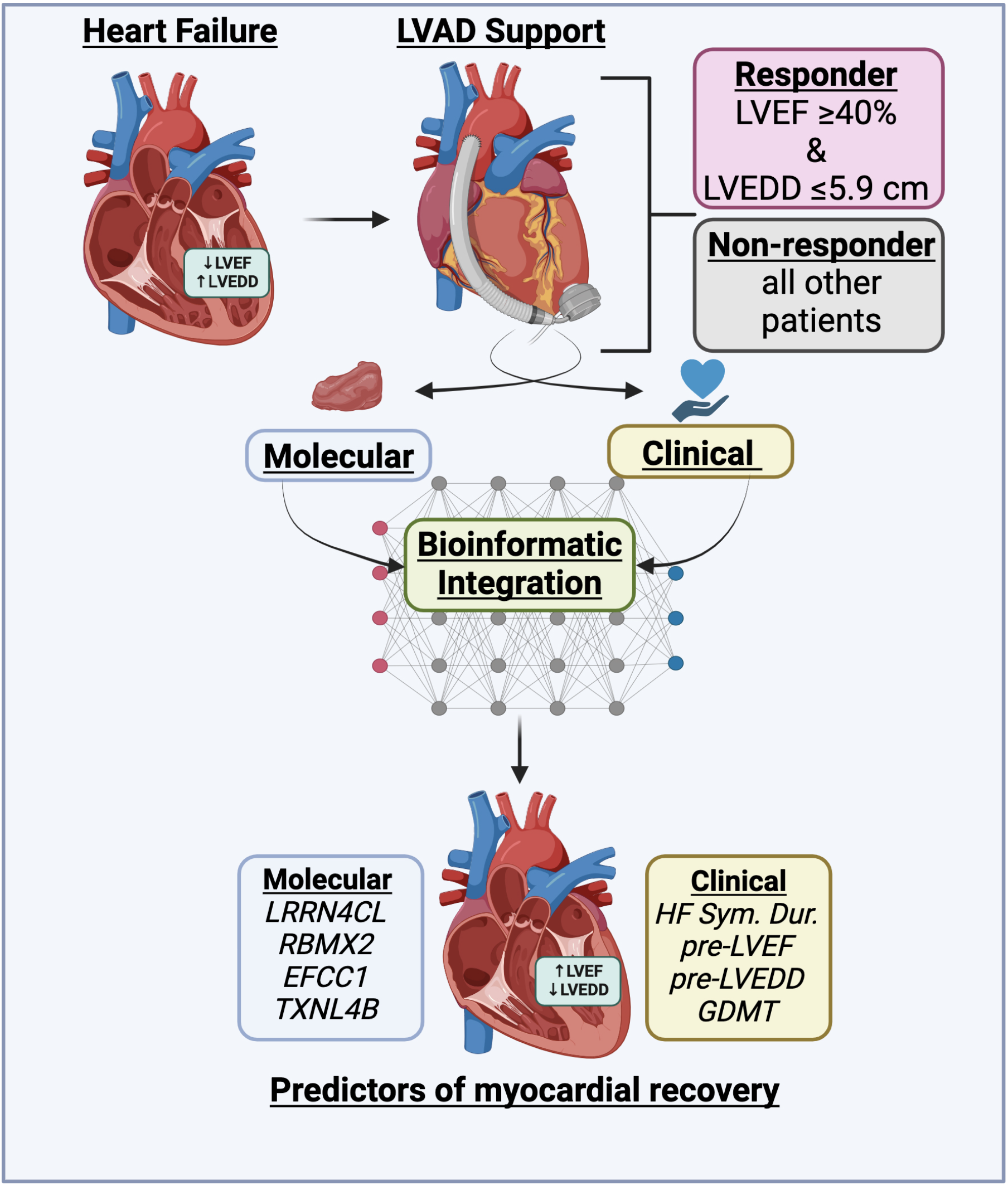

## INTRODUCTION

Heart failure (HF) is a progressive, chronic disease associated with high morbidity and mortality (1). Left ventricular assist devices (LVADs) are an established therapeutic approach for advanced HF patients with refractory HF symptoms despite guideline-directed medical therapy and are increasingly used either as a bridge to heart transplantation, or as destination lifetime therapy (2). The placement of an LVAD offers a unique investigational model by providing left ventricular (LV) apical myocardial tissue at the time of implantation. Previously it has been shown that LVADs, through volume and pressure unloading of the failing LV, can enable structural and functional cardiac improvement (3–9). On the molecular level, LVADs result in positive adaptations to cardiomyocyte metabolism, excitation-contraction coupling, and gene expression (10–15). Although certain clinical characteristics (6, 8, 9, 16) and biological observations (12, 14, 17, 18) have been associated with a higher tendency for structural and functional cardiac improvement, the interplay between clinical outcomes and underlying biological phenomena driving these changes is not well-understood. Currently, there is a paucity of accurate predictive markers and molecular mechanism to forecast LVAD-mediated myocardial recovery among advanced HF patients and innovative bioinformatic approaches are needed (19, 20).

Analyses that combine clinical *and* molecular profiles using machine learning (ML) has proven useful for the diagnosis, prognosis, and classification of other human diseases (21–23), but to our knowledge has not been used to derive predictors of myocardial recovery. Compared to traditional pre-specified regression-based models, ML has the capacity to uncover complex interactions between variables, particularly for molecular-associated phenotypes that exist across multiple biological pathways, such as HF. Our study used supervised ML to integrate clinical variables and transcriptomics pre-LVAD implantation which were associated with structural and/or functional cardiac improvement following LVAD unloading. We hypothesized that this approach would predict novel molecular markers and will improve the “out-of-sample” prediction associated with LVAD-mediated myocardial recovery *in-silico* (24). By doing so, this investigation aims to elucidate clinical and molecular predictors of LVAD-mediated myocardial recovery that could lead to biological understanding of myocardial recovery and novel therapeutic targets in LVAD recipients and the broader HF population.

## METHODS

The data that support the findings of this study are available from the corresponding author upon reasonable request, provided the request made does not violate human subject confidentiality protocols.

### Study Population

Advanced HF patients receiving continuous-flow LVADs across five medical centers in the United States comprised our study cohort. Patients were prospectively enrolled at the institutions of the Utah Cardiac Recovery (UCAR) Program (University of Utah Health & School of Medicine, Intermountain Medical Center, and Salt Lake City VA Medical Center), located in Salt Lake City, UT, and retrospectively reviewed at Allegheny General Hospital (Pittsburgh, PA), and the University of Louisville (Louisville, KY).

Patients who required LVAD support due to acute causes of HF (acute myocardial infarction, acute myocarditis, post-cardiac surgery cardiogenic shock, etc.) including those with normal LV chamber size were excluded, as they may be considered having a different underlying pathophysiology, rendering them more prone to myocardial recovery. Subjects were also excluded if previously diagnosed with hypertrophic or infiltrative cardiomyopathies, had a baseline LV ejection fraction (LVEF) ≥40%, withdrew consent, or had inadequate (< 3 months) post-LVAD follow up (i.e., early heart transplantation, early death, or loss to follow-up).

### Clinical Protocol

Data collection included demographics, duration of HF symptoms prior to LVAD implant, co-morbidities, medications, laboratories, hemodynamics, and cardiac imaging information. Hemodynamic data were obtained during the right heart catheterization prior to and closest to LVAD implantation, while imaging and laboratory data were collected within one week prior to implantation and stored for offline analysis.

After LVAD implantation, patients were medically managed at the discretion of the treating physicians within the participating institutions per established standard HF therapy guidelines. The effect of LVAD unloading on cardiac size, shape, and function was assessed by echocardiography (see below) and invasive hemodynamic measurements. LVAD speed was adjusted to optimize flow and LV decompression with positioning of the interventricular and interatrial septa in the midline, minimal mitral valve regurgitation, and intermittent aortic valve opening, in order of decreasing priority. Speed adjustments were made as indicated by patients’ symptoms and/or clinical events.

### Echocardiography

Echocardiographic examinations were performed at the echocardiography laboratories of the participating institutions and were digitally stored. At the institutions comprising the UCAR Program the echocardiograms were performed within 1-month before LVAD implantation, and then at 1, 2, 3, 6, 9, and 12 months on LVAD support. At Allegheny General Hospital and the University of Louisville post-LVAD echocardiograms were performed at 1, 3, 6, and 12 months following LVAD implantation. Guidelines of the American Society of Echocardiography and the European Association of Cardiovascular Imaging were followed and included two-dimensional, M-mode, and Doppler modalities (25). Patients responding favorably to LVAD support were defined as

- *Responders (R)*: with a follow-up LVEF ≥40 %, and LV end-diastolic diameter (LVEDD) ≤5.9 cm within the year post-LVAD implantation (8)
- *Non-responders (NR)*: remaining patients who did not meet criteria to be classified as responders

Patients undergoing mechanical unloading may experience structural and/or functional myocardial improvement to varying degrees (8). Moreover, patients consistently show structural cardiac improvement, while fewer experience functional improvement (6, 8). To independently examine functional and structural myocardial improvement, we evaluated improvements in LVEF and LVEDD within the year post-LVAD implantation as continuous variables.

### Human Myocardial Tissue Procurement

Human myocardial tissue was prospectively collected from the LV apical core at the time of LVAD implantation. Myocardial tissue was also procured from non-failing donor hearts that were ineligible for transplantation due to non-cardiac reasons (infection, donor-recipient size mismatch, etc.). Predictions, gene expression, and protein abundance were assessed using *only tissue collected at the pre-timepoint*. Tissue was immediately frozen in liquid nitrogen before storing it at -80°C.

### RNA Isolation, Sequencing and Bioinformatics

Total RNA was isolated from 50 mg of frozen human myocardial LV tissue samples using the miRNeasy Mini Kit (Qiagen: Hilden, DE). To remove all DNA, an on-column RNase-free deoxyribonuclease digestion was used. All samples were processed at the same time for dispersion estimation and normalization, with contrasts applied to specific condition groups for differential expression testing. Samples from different centers were processed at one location at the Genomic Core facility of the Huntsman Cancer Institute at the University of Utah. A batch factor was introduced in the model to account for different sequencing batches. Differentially expressed genes were filtered using the criteria: adjusted *p-value <0.05*, absolute log^2^ fold change >0.585 (or linear fold change >1.5), and a normalized base mean count of 30 (to remove any significant genes with very low expression). Bioinformatic analysis of transcripts predicted to influence LVAD-mediated myocardial recovery was performed using the Ingenuity Pathway Analysis (IPA) and STRING database for protein-protein interactions. Using the ΔΔCT method, quantitative real-time polymerase chain reaction (RT-qPCR) confirmed relative expression levels (normalized to vinculin) of predicted transcripts associated with LVAD-mediated myocardial recovery, improved LVEF and LVEDD, at the pre-LVAD timepoints.

### Data Processing

Predictions depended on feature selection *at the pre-LVAD timepoint* from a pool of 22,373 mRNA transcripts and 59 clinical variables. We removed clinical variables with at least 40% missing data overall from inclusion in the analysis. This resulted in 51 candidate clinical features from five individual hospitals. For the remaining clinical variables, if there were any missing data, we used multiple imputation with chained equations, creating five complete data sets using the mice package in R-software (26). Multiple imputation ensures valid estimation and inference under the missing-at-random assumption. For this investigation we were only interested in measuring feature importance and measuring performance of a predictive model, so we stacked our five complete data sets to account for the variation from imputing the missing data as opposed to pooling results (27).

### Predictive Models and Assessing Performance

We conducted 100 iterations of repeated cross-validation where, in each iteration, we randomly selected 90% of our patients as our “training dataset” with equal representation from each hospital and LVAD response (R or NR) and used the remaining patients as a “testing dataset” to assess the performance of each model. Given the small sample size and lack of external testing set, we trained on 90% of our data in order to maximize the generalizability of the cross-validation estimate of performance to predictions made on new data with a model which would be trained on our full data. For the prediction of LVAD response (binary outcome), we assessed the discriminatory performance of the model using the area under the receiver operating characteristic curve (AUC: higher value represents better performance) and assessed whether it achieved weak calibration by estimating the calibration intercept and slope (28). Given the imbalance of the binary outcome in our data, we additionally present the area under the precision-recall curve (PRAUC), the combination of precision (positive predictive value) and recall (sensitivity) for the models with the best discriminatory performance. For the prediction of LVEF and LVEDD (continuous outcomes), we assessed performance using mean absolute error (MAE: lower value represents better performance) (29). Then, we averaged each patient’s predictions/imputations to visualize patient-specific error.

Both within cross-validation and on the entire dataset, we conducted feature selection to order features based on their importance to build a parsimonious model and to assess overfitting. Due to the large number of variables, we conducted screening of the mRNA transcripts in two stages to limit the number of features considered in each model: 1) traditional screening (simple linear or logistic regression) and then 2) ML-based screening, using the random forest conditional permutation algorithm. Using this approach, we tested models of varying parameter subset sizes as well as models with only features from clinical data, only mRNA transcript data, and their combination. Further details regarding the screening methods can be viewed in Figure 1. Additionally, we present partial dependency plots of the top 10 clinical variables and the top 10 mRNA variables for each outcome resulting from a random forest model fit on the full dataset using these variable sets. Last, the performance of our predictive model was compared to existing LVAD-mediated myocardial recovery predictive models.

**Figure 1.**
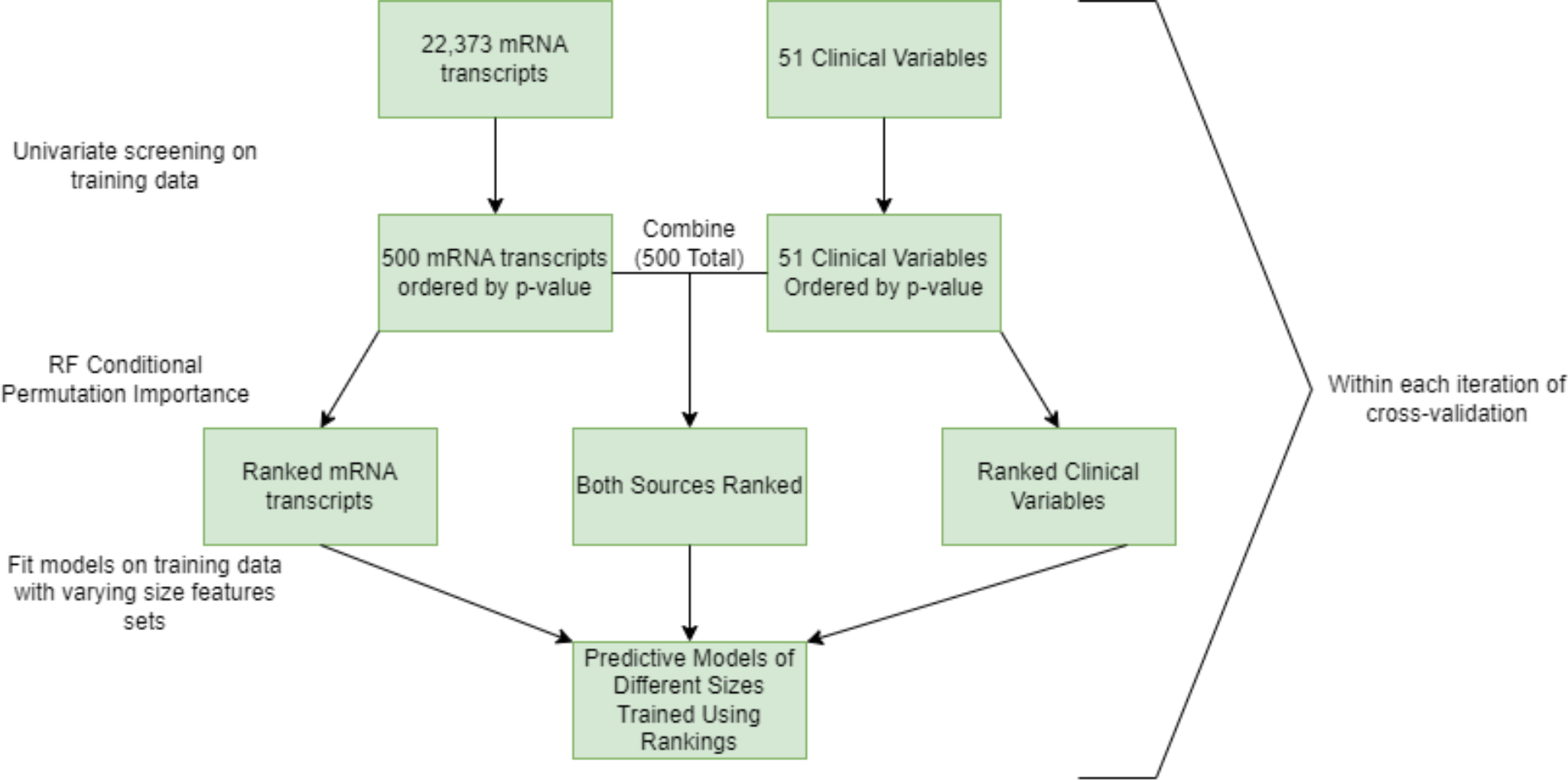
Screening and Testing Methodology for the mRNA transcripts and clinical variables.

All analyses were conducted in R 4.1.0 (R Core Team, 2014). We fit the predictive models using the following functions: glm (logistic regression), cforest (random forest, 5000 trees), gbm (gradient boosted regression), svm (support-vector machine), each using default settings unless otherwise specified (30).

### Statistics

The Shapiro-Wilk and D’Agostino-Pearson omnibus tests were used to verify the normality of data. ROUT testing (GraphPad Prism v9.01, San Diego, CA) was used to identify any significant outliers, and if identified were removed prior to analysis. If data were not normally distributed, a logarithmic transformation was performed, following which, a one-way ANOVA was used to compare the relative gene expression in the following groups: 1) Donors, 2) Non-Responders (NR), and 3) Responders (R). An α level of *p* < 0.05 was set *a priori,* and if necessary, a Tukey’s honest significant difference post-hoc test was used for multiple comparisons.

### Study Approval

At all centers, the study was approved by institutional review board committees and subjects provided written informed consent for participation in the study. Human myocardial tissue was collected under approved IRB protocols.

## RESULTS

### Patient Characteristics

Our study cohort comprised 208 chronic advanced HF patients undergoing durable, continuous-flow LVAD implantation in the following institutions (UCAR Program; n=123, Allegheny General Hospital; n=53, University of Louisville; n=32, Supplemental Table 1). Baseline clinical characteristics of R and NR can be seen in Table 1. Overall, the R group were younger, had less severe HF symptoms for a shorter time (NYHA Classification), were less commonly on cardiac resynchronization therapy or an implantable cardioverter defibrillator, and had a less dilated LV prior to LVAD implant, when compared to NR.

**Table 1:**
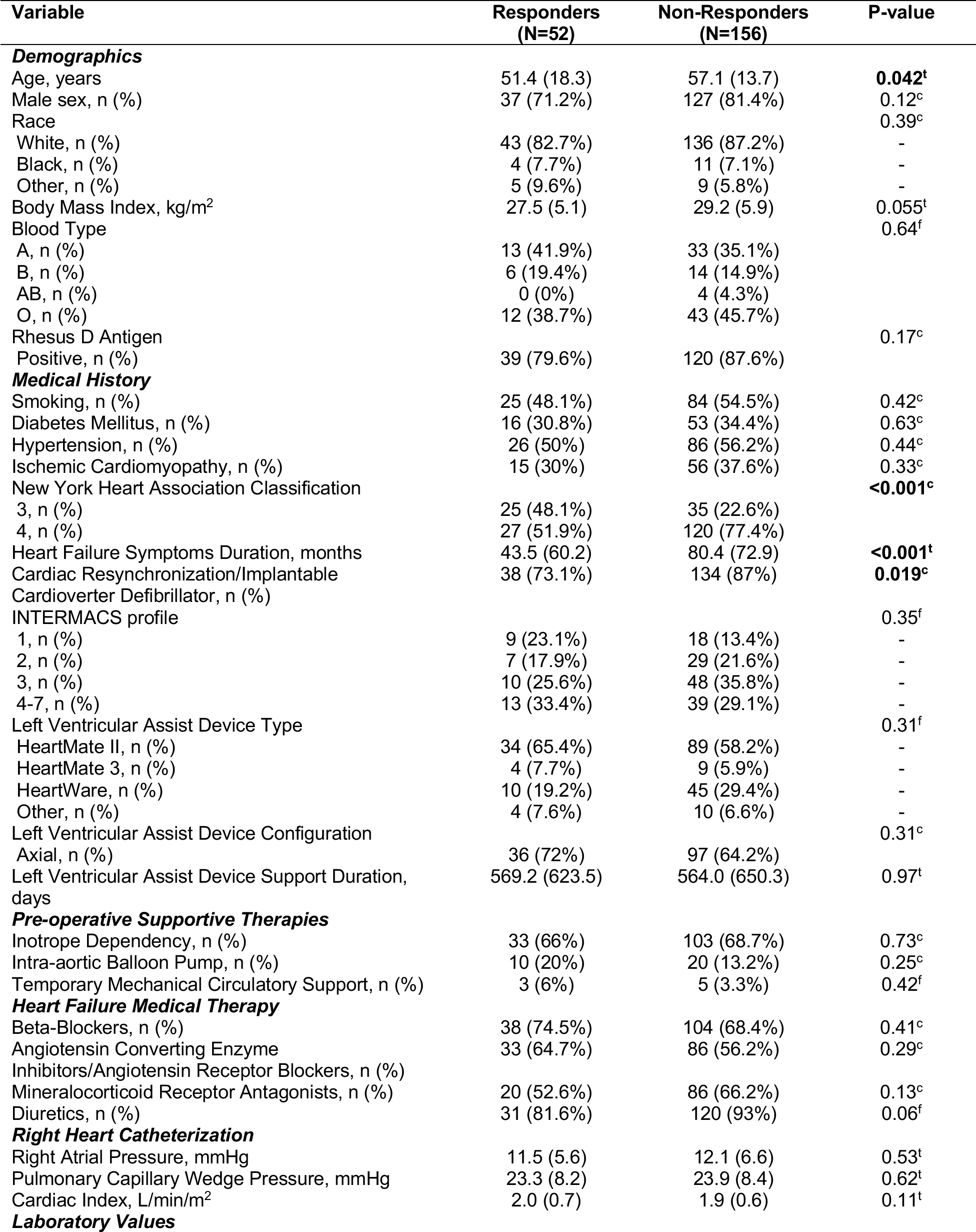
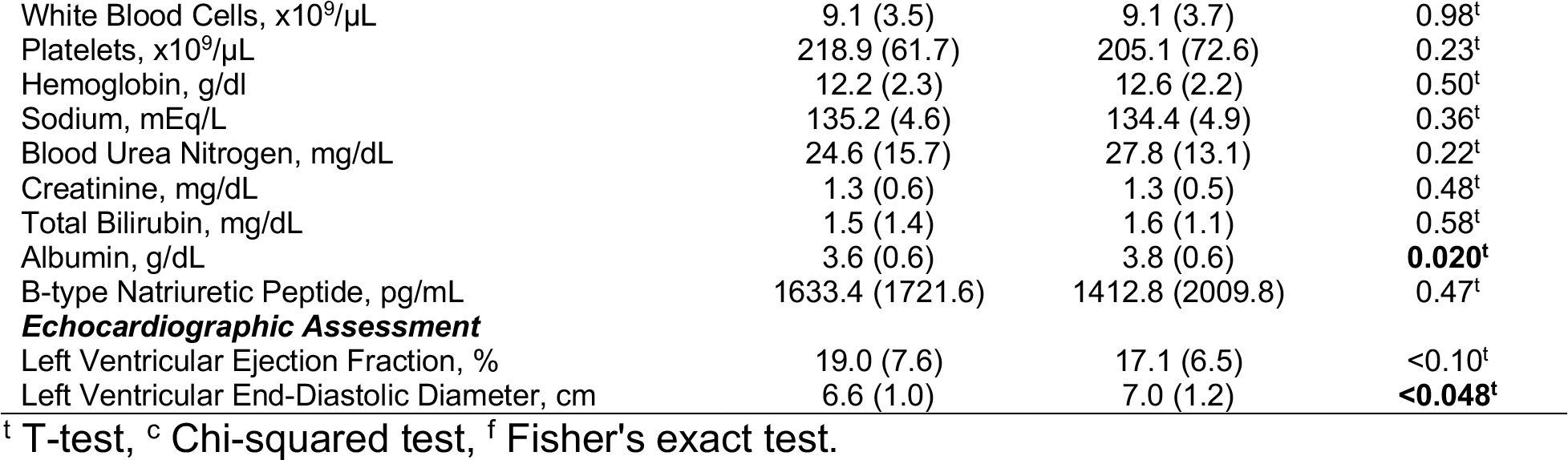
Baseline clinical characteristics of LVAD responders and non-responders.

| Variable | Responders<br>(N=52) | Non-Responders<br>(N=156) | P-value |
| --- | --- | --- | --- |
| **Demographics** |  |  |  |
| Age, years | 51.4 (18.3) | 57.1 (13.7) | **0.042<sup>t</sup>** |
| Male sex, n (%) | 37 (71.2%) | 127 (81.4%) | 0.12<sup>c</sup> |
| Race |  |  | 0.39<sup>c</sup> |
| White, n (%) | 43 (82.7%) | 136 (87.2%) | - |
| Black, n (%) | 4 (7.7%) | 11 (7.1%) | - |
| Other, n (%) | 5 (9.6%) | 9 (5.8%) | - |
| Body Mass Index, kg/m<sup>2</sup> | 27.5 (5.1) | 29.2 (5.9) | 0.055<sup>t</sup> |
| Blood Type |  |  | 0.64<sup>f</sup> |
| A, n (%) | 13 (41.9%) | 33 (35.1%) |  |
| B, n (%) | 6 (19.4%) | 14 (14.9%) |  |
| AB, n (%) | 0 (0%) | 4 (4.3%) |  |
| O, n (%) | 12 (38.7%) | 43 (45.7%) |  |
| Rhesus D Antigen |  |  | 0.17<sup>c</sup> |
| Positive, n (%) | 39 (79.6%) | 120 (87.6%) |  |
| **Medical History** |  |  |  |
| Smoking, n (%) | 25 (48.1%) | 84 (54.5%) | 0.42<sup>c</sup> |
| Diabetes Mellitus, n (%) | 16 (30.8%) | 53 (34.4%) | 0.63<sup>c</sup> |
| Hypertension, n (%) | 26 (50%) | 86 (56.2%) | 0.44<sup>c</sup> |
| Ischemic Cardiomyopathy, n (%) | 15 (30%) | 56 (37.6%) | 0.33<sup>c</sup> |
| New York Heart Association Classification |  |  | **<0.001<sup>c</sup>** |
| 3, n (%) | 25 (48.1%) | 35 (22.6%) |  |
| 4, n (%) | 27 (51.9%) | 120 (77.4%) |  |
| Heart Failure Symptoms Duration, months | 43.5 (60.2) | 80.4 (72.9) | **<0.001<sup>t</sup>** |
| Cardiac Resynchronization/Implantable | 38 (73.1%) | 134 (87%) | **0.019<sup>c</sup>** |
| Cardioverter Defibrillator, n (%) |  |  |  |
| INTERMACS profile |  |  | 0.35<sup>f</sup> |
| 1, n (%) | 9 (23.1%) | 18 (13.4%) | - |
| 2, n (%) | 7 (17.9%) | 29 (21.6%) | - |
| 3, n (%) | 10 (25.6%) | 48 (35.8%) | - |
| 4-7, n (%) | 13 (33.4%) | 39 (29.1%) | - |
| Left Ventricular Assist Device Type |  |  | 0.31<sup>f</sup> |
| HeartMate II, n (%) | 34 (65.4%) | 89 (58.2%) | - |
| HeartMate 3, n (%) | 4 (7.7%) | 9 (5.9%) | - |
| HeartWare, n (%) | 10 (19.2%) | 45 (29.4%) | - |
| Other, n (%) | 4 (7.6%) | 10 (6.6%) | - |
| Left Ventricular Assist Device Configuration |  |  | 0.31<sup>c</sup> |
| Axial, n (%) | 36 (72%) | 97 (64.2%) |  |
| Left Ventricular Assist Device Support Duration, days | 569.2 (623.5) | 564.0 (650.3) | 0.97<sup>t</sup> |
| **Pre-operative Supportive Therapies** |  |  |  |
| Inotrope Dependency, n (%) | 33 (66%) | 103 (68.7%) | 0.73<sup>c</sup> |
| Intra-aortic Balloon Pump, n (%) | 10 (20%) | 20 (13.2%) | 0.25<sup>c</sup> |
| Temporary Mechanical Circulatory Support, n (%) | 3 (6%) | 5 (3.3%) | 0.42<sup>f</sup> |
| **Heart Failure Medical Therapy** |  |  |  |
| Beta-Blockers, n (%) | 38 (74.5%) | 104 (68.4%) | 0.41<sup>c</sup> |
| Angiotensin Converting Enzyme | 33 (64.7%) | 86 (56.2%) | 0.29<sup>c</sup> |
| Inhibitors/Angiotensin Receptor Blockers, n (%) |  |  |  |
| Mineralocorticoid Receptor Antagonists, n (%) | 20 (52.6%) | 86 (66.2%) | 0.13<sup>c</sup> |
| Diuretics, n (%) | 31 (81.6%) | 120 (93%) | 0.06<sup>f</sup> |
| **Right Heart Catheterization** |  |  |  |
| Right Atrial Pressure, mmHg | 11.5 (5.6) | 12.1 (6.6) | 0.53<sup>t</sup> |
| Pulmonary Capillary Wedge Pressure, mmHg | 23.3 (8.2) | 23.9 (8.4) | 0.62<sup>t</sup> |
| Cardiac Index, L/min/m<sup>2</sup> | 2.0 (0.7) | 1.9 (0.6) | 0.11<sup>t</sup> |
| **Laboratory Values** |  |  |  |
| White Blood Cells, x10<sup>9</sup>/μL | 9.1 (3.5) | 9.1 (3.7) | 0.98<sup>t</sup> |
| Platelets, x10<sup>9</sup>/μL | 218.9 (61.7) | 205.1 (72.6) | 0.23<sup>t</sup> |
| Hemoglobin, g/dl | 12.2 (2.3) | 12.6 (2.2) | 0.50<sup>t</sup> |
| Sodium, mEq/L | 135.2 (4.6) | 134.4 (4.9) | 0.36<sup>t</sup> |
| Blood Urea Nitrogen, mg/dL | 24.6 (15.7) | 27.8 (13.1) | 0.22<sup>t</sup> |
| Creatinine, mg/dL | 1.3 (0.6) | 1.3 (0.5) | 0.48<sup>t</sup> |
| Total Bilirubin, mg/dL | 1.5 (1.4) | 1.6 (1.1) | 0.58<sup>t</sup> |
| Albumin, g/dL | 3.6 (0.6) | 3.8 (0.6) | **0.020<sup>t</sup>** |
| B-type Natriuretic Peptide, pg/mL | 1633.4 (1721.6) | 1412.8 (2009.8) | 0.47<sup>t</sup> |
| ***Echocardiographic Assessment*** |  |  |  |
| Left Ventricular Ejection Fraction, % | 19.0 (7.6) | 17.1 (6.5) | <0.10<sup>t</sup> |
| Left Ventricular End-Diastolic Diameter, cm | 6.6 (1.0) | 7.0 (1.2) | **<0.048<sup>t</sup>** |

<sup>t</sup> T-test, <sup>c</sup> Chi-squared test, <sup>f</sup> Fisher's exact test.

### Molecular and Clinical Variables Accompanying Myocardial Recovery

Cross-validation results for predicting LVAD-mediated myocardial recovery showed that using both data sources in the gradient boosted machine (gbm) model with 80 features achieves an AUC of 0.731±0.15 (Figure 2A). However, the PRAUC is maximized under the gbm model with only 30 features (PRAUC = 0.445±0.253), outperforming the 80-variable model by 2% (PRAUC = 0.425±0.242). Using only clinical variables, the random forest (rf) with 20 variables achieved an AUC of 0.727±0.14 and a PRAUC of 0.471±0.237 in cross-validation (Figure 2B). Using transcriptomics alone, the gbm model with 70 features achieves an AUC of 0.713±0.14 and a PRAUC of 0.388±0.227 in cross-validation (Figure 2C). As a comparison to the previously reported Interagency Registry for Mechanically Assisted Circulatory Support (INTERMACS) Cardiac Recovery (ICAR) score predicting patients achieving significant functional and structural cardiac improvement and subsequent LVAD removal, we fit a logistic regression model with only the variables employed in the ICAR score (9), which resulted in an AUC of 0.735±0.134 (Figure 2D). Using a paired randomization test, we compared the average AUC from the ICAR score with both the 80 variable gbm model using both data sources and the 20 variable rf model with the clinical data source and found no evidence of a difference in average AUC between them (p-value =0.80 and p-value=0.49, respectively).

**Figure 2.**
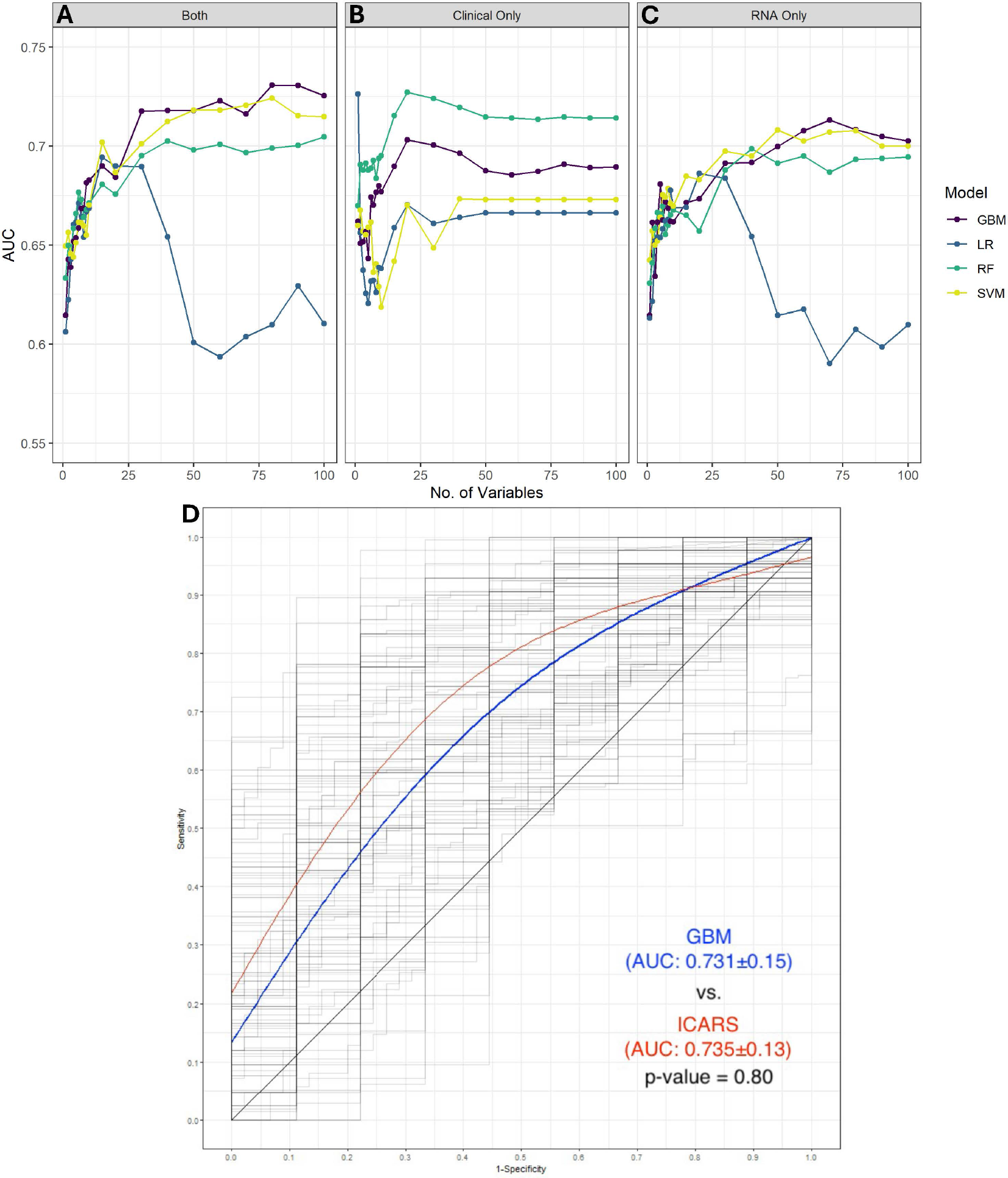
Cross-validated AUCs by both clinical and mRNA variables (**A**), only clinical variables (**B**), or only mRNA variables (**C**) and the number of variables used in the model for predicting LVAD-mediated myocardial recovery. Variables were ranked and selected using a two-stage screening process. **D**) A smooth averaging of the ROC curves for the GBM model (solid blue line) with 80 variables, and the ICAR score (solid red line) using cubic smoothing splines with 4 degrees of freedom as well as the ROC curve from each of the 100 iterations of cross-validation (shadowy lines). Gradient Boosted Regression (gbm: 5000 trees), Logistic Regression (glm), Random Forest (rf: 5000 trees), and Support Vector Machine (svm: polynomial regression). A paired randomization test was used to compare GBM against ICARS resulting in a p-value of 0.80.

We assessed the calibration of the candidate models and found the calibration of the 80 variable gbm model employing both clinical and transcriptomic data to be substandard. However, the rf model with 20 variables using only clinical data had cross-validated averages for the calibration intercept of -0.46 (95% CI: -1.10, 0.15), with zero as the target and slope of 1.16 (95% CI: 0.19, 2.14) with one as the target. There is evidence this model achieves weak calibration, though the point estimates suggest predictions were systematically extreme compared to the observed proportion (Figure 3). Using the entire dataset, we found that HF symptoms duration, use of mineralocorticoid receptor antagonists (MRA), and LVAD configuration (axial vs. centrifugal) were the top-ranking clinical variables for predicting R (Table 2). Specifically, the partial dependency plots (Supplemental Figure 1) show that patients with shorter HF symptoms duration, who had not been on MRA therapy, and were supported with an axial-flow LVAD had a higher probability of meeting responders’ criteria. Additionally, the top-ranking mRNA transcripts for the prediction of LVAD-mediated myocardial recovery were cell cycle regulator of non**-**homologous end-joining recombination (*CYREN*), activating transcription factor 3 (*ATF3*), dual specificity phosphatase 5 (*DUSP5*), and leucine-rich repeat neuronal 4C like (*LRRN4CL*). Higher expression levels of *CYREN* but lower expression levels of *ATF3*, *DUSP5*, and *LRRN4CL* were associated with an increased probability of myocardial recovery.

**Figure 3:**
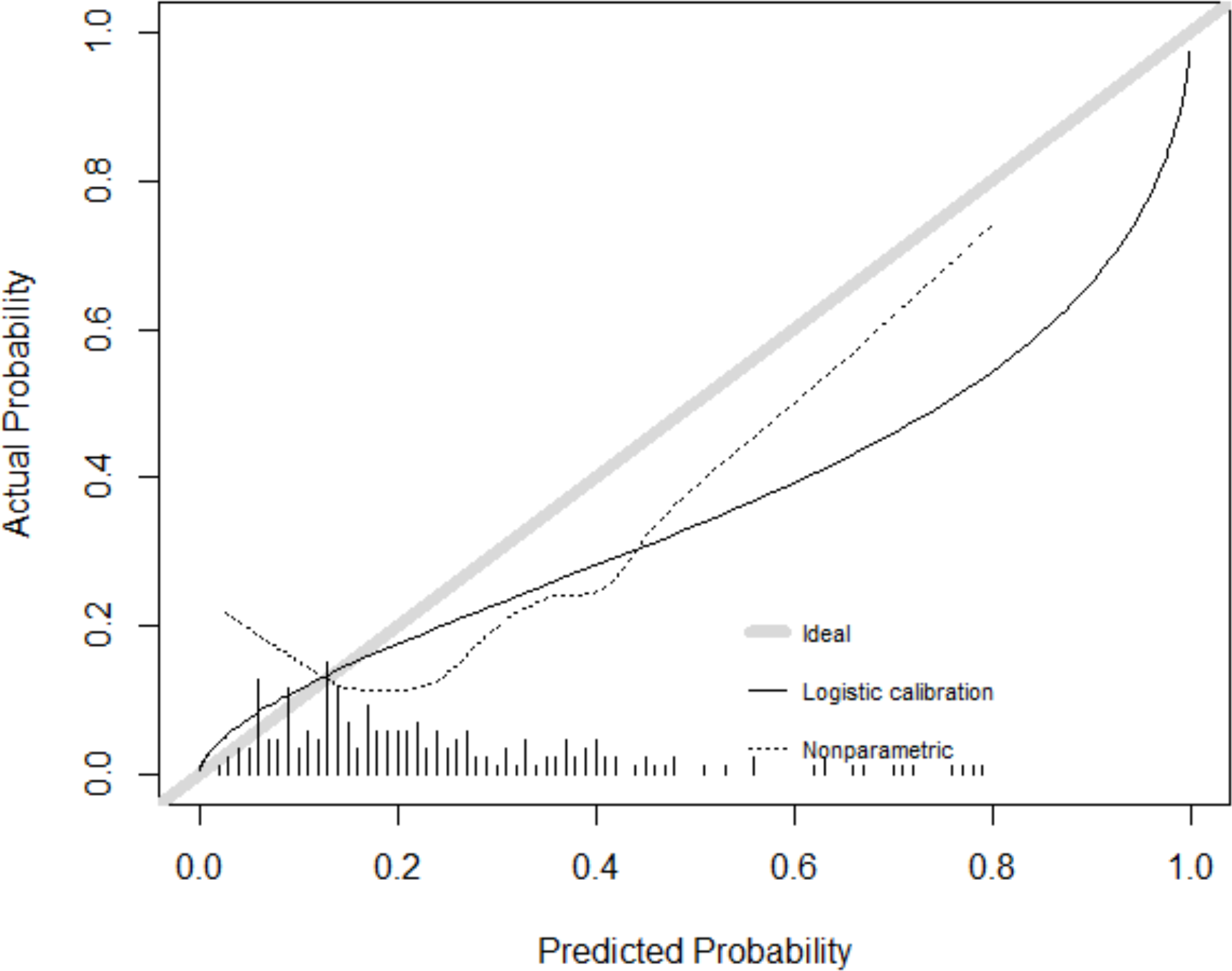
Calibration of RF model with clinical data (20 variables). Cross-validated averages were obtained for the calibration intercept of -0.46 (95% CI: -1.10, 0.15), with 0 as the target and slope of 1.16 (95% CI: 0.19, 2.14) with 1 as the target. The calibration plot shows the average predicted values from cross-validation (solid and dotted) were systematically greater than the ideal value (grey).

**Table 2:** Variable importance for predicting LVAD-mediated myocardial recovery.

| *Rank* | *Clinical* | *Mean*<br>*Decrease*<br>*AUC* | *Gene Code* | *Mean*<br>*Decrease*<br>*AUC* |
| --- | --- | --- | --- | --- |
| 1 | HF Symptoms Duration | 0.071 | CYREN | 0.012 |
| 2 | Mineralocorticoid Receptor<br>Antagonists | 0.015 | ATF3 | 0.012 |
| 3 | LVAD configuration (axial<br>vs. centrifugal) | 0.015 | DUSP5 | 0.011 |
| 4 | LVAD Type | 0.013 | **LRRN4CL** | 0.009 |
| 5 | LVEF | 0.012 | APRT | 0.009 |
| 6 | History of Hypertension | 0.012 | TRDMT1 | 0.007 |
| 7 | Creatinine | 0.010 | HBB | 0.007 |
| 8 | Body Mass Index | 0.010 | LONRF1 | 0.006 |
| 9 | LVEDD | 0.009 | MORF4L1P1 | 0.006 |
| 10 | Rhesus D Antigen | 0.009 | ZFTA | 0.005 |

Clinical & RNA variables from random forests in the 2-stage screening process were used. The variable importance is ranked 1-10. AUC; area under curve, HF; heart failure, LVAD; left ventricular assist device, LVEDD; left ventricular end-diastolic diameter, LVEF; left ventricular ejection fraction

The top ten predictive transcripts were further analyzed via IPA and STRING bioinformatics software. IPA showed the coronavirus pathogenesis pathway (p=0.0039) as the top canonical pathway associated with the predictive transcripts for LVAD-mediated myocardial recovery (Figure 4A). The STRING protein-protein interactions showed nodes positioned around *ATF3*, *HBB*, and *APRT* which have been enriched with other potential interactors such as *SMAD3*, *HBA1*, and *ADK* (Supplemental Figure 2A). To further assess variable importance, we performed a univariate logistic regression analysis with each of the clinical and RNA variables and independent variables. HF symptoms duration (Figure 4B), *LRRN4CL*, and *ATF3* (Figure 4C-D) appeared as significant (p<0.05) clinical and biological variables that were also top variables from the random forest importance metrics.

**Figure 4:**
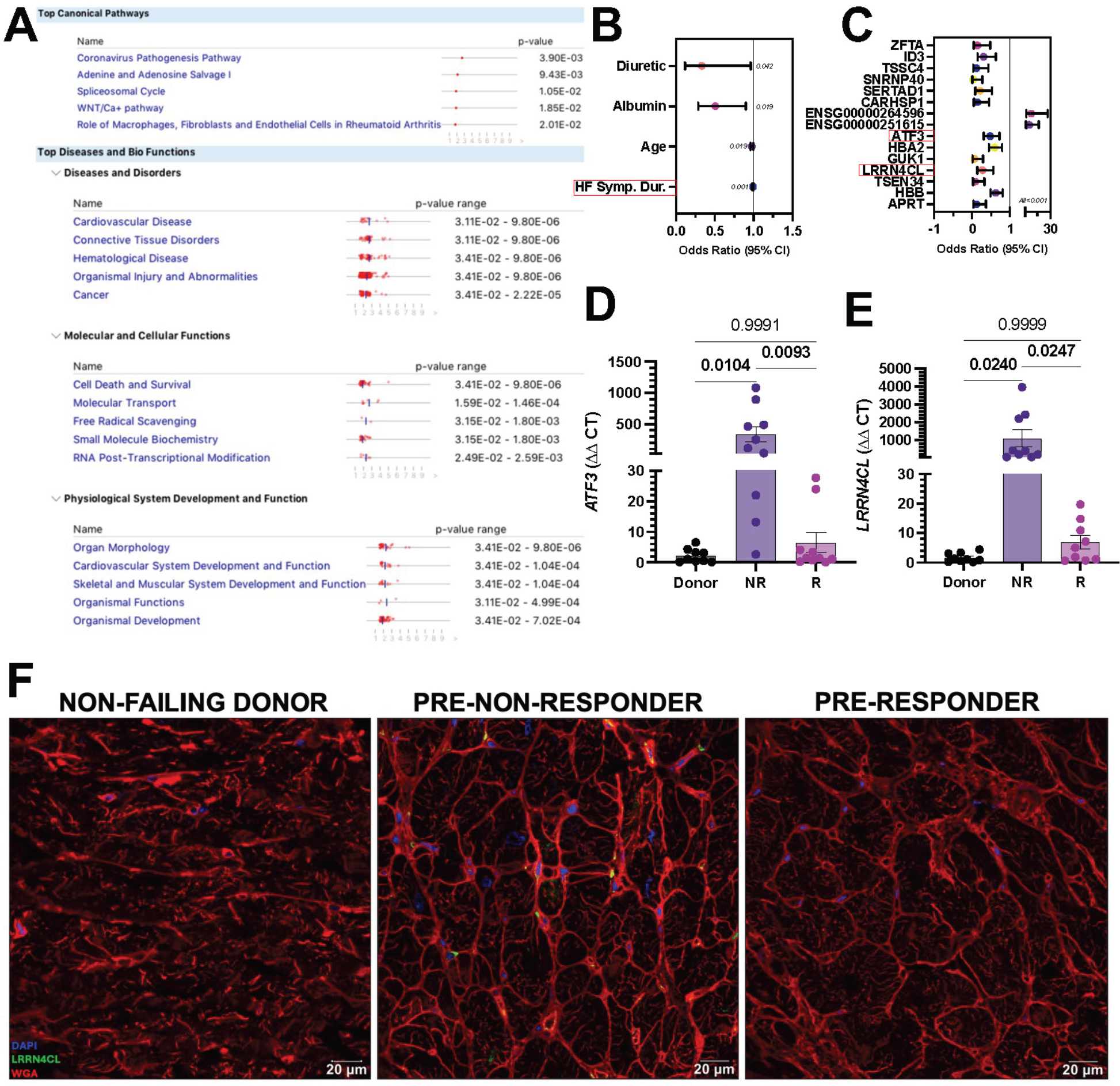
Bioinformatics analysis for the top 10 predictive RNA transcripts associated with LVAD-mediated myocardial recovery. **A)** Ingenuity Pathway Analysis shows the top canonical pathways, diseases, and bio-functions associated with the transcripts for recovery. **B-C)** Odds ratios for the significantly identified clinical variables and mRNA transcripts using logistic regression analysis. Red boxes indicate variables that were identified via ML. **D-E)** Relative gene expression of *ATF3* and *LRRN4CL* as determined by RT-qPCR. **F)** Immunofluorescent confocal microscopy of DAPI (blue), WGA (red), and LRRN4CL (green) at 60X magnification. Donor (non-failing human hearts), non-responders (NR), responders (R).

*LRRN4CL* emerged from bioinformatic integration as the top predictor of myocardial recovery, improved cardiac function and structure (see below). Thus, to provide mechanistic insights associated with myocardial improvement between R and NR, we sought to characterize *LRRN4CL* more in-depth. The *LRRN4CL* transcriptional signature presents with a higher expression in the cardiac tissue of NR compared to R and non-failing donors before LVAD implant. RT-qPCR shows *LRRN4CL* to be dysregulated in NR and normalized in R compared to non-failing donors (Figure 4E). From cellular indexing of transcriptomes and epitopes by sequencing (CITE-seq) (31), the expression of *LRRN4CL* is expressed in fibroblasts and lymphocytes (Supplemental Figure 2B). Next, confocal immunofluorescence microscopy using human cardiac tissue show increased fluorescence of LRRN4CL in the extracellular space of NR compared with R and non-failing donors (Figure 4F). Last, we probed for LRRN4CL using western blotting and showed the relative protein abundance was increased in the NR when compared only with R (Supplemental Figure 2C). Taken together, these data indicate the impact of *LRRN4CL* expression in HF and myocardial recovery.

### Molecular and Clinical Data Before LVAD Associated with Functional or Structural Cardiac Improvement

Patients undergoing mechanical unloading with LVAD support may experience structural and/or functional myocardial improvement (4–7, 9) to varying degrees (8). In fact, these patients consistently show structural cardiac improvement, while a lower percentage experiences functional improvement (6, 8). Furthermore, LVAD-mediated myocardial recovery may represent a spectrum of response and not a binary outcome. With this in mind, we also attempted to predict improvements to LVEF and LVEDD independently, as continuous variables.

### Predicting Functional Cardiac Improvement

Cross-validation revealed that the 10-variable rf model only using clinical variables had the highest performance in predicting LVEF improvement with an MAE of 9.93±1.71 on average (Figure 5). The svm model using both data sources achieved an MAE of 10.3±1.43 on average with only one variable. When using transcriptomics, the 70-variable rf model achieved an MAE of 11.2±1.77 on average.

**Figure 5:**
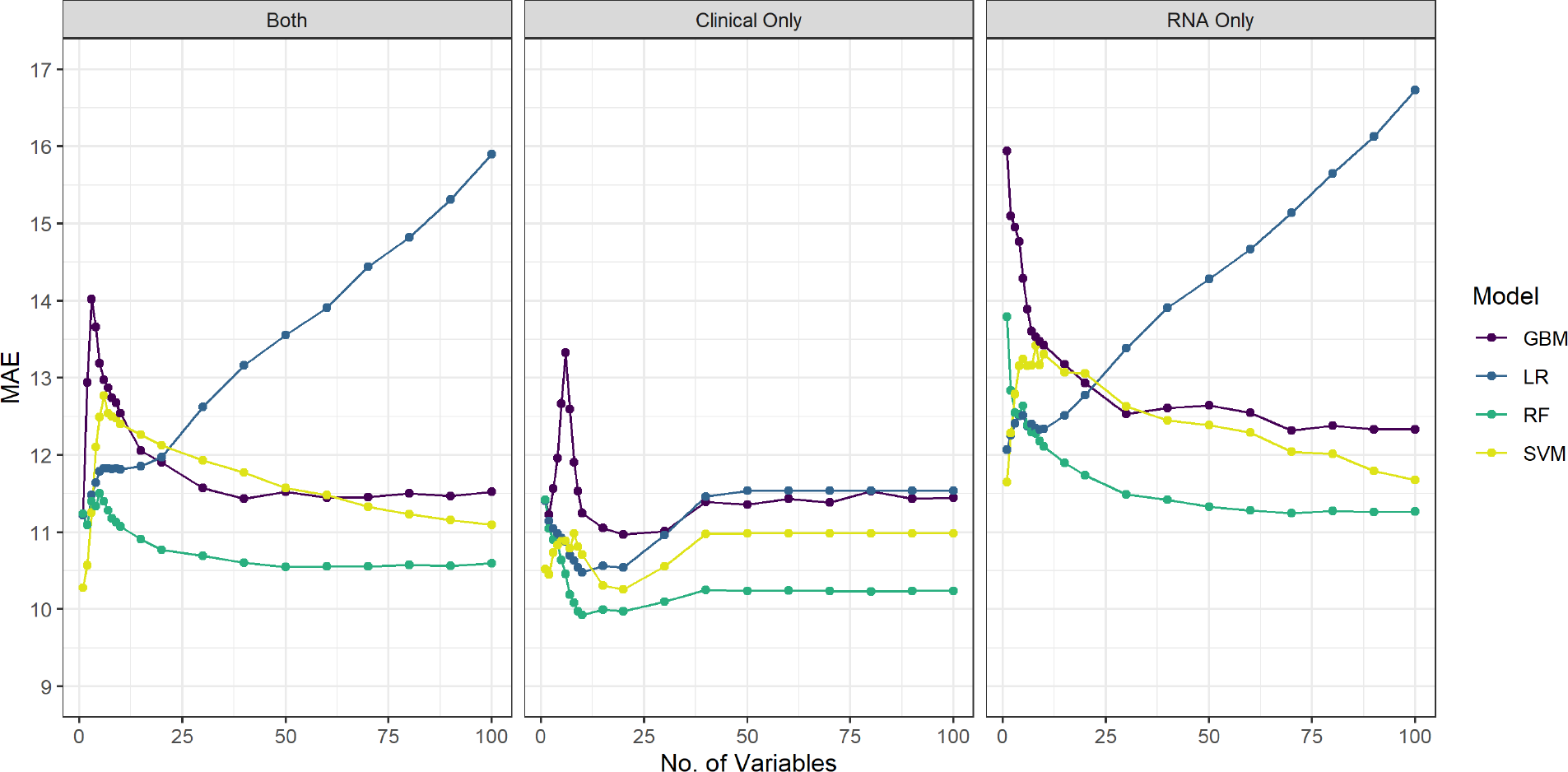
Cross-validated Mean Absolute Error (MAE) by both data sources (left), clinical characteristics (middle) or mRNA transcripts (right) and number of variables used in the model for predicting improved myocardial function (LVEF). A lower MAE indicates better predictive performance. Variables were selected using a two-stage screening process. Gradient Boosted Regression (gbm: 5000 trees), Logistic Regression (glm), Random Forest (rf: 5000 trees), and Support Vector Machine (svm: polynomial regression).

Baseline LVEF, post-LVAD LVEF, and the predicted post-LVAD LVEF values for each patient show our prediction is conservative for patients with the greatest improvement (Left; Supplemental Figure 3). Our prediction error increases as we move away from the average post-LVAD LVEF (Right; Supplemental Figure 3).

We found that sex, LVEDD, and LVAD configuration (axial vs. centrifugal) were the most important clinical variables for predicting LVEF improvement post-LVAD when using the training dataset (Table 3). The partial dependency plots for each variable are shown in Supplemental Figure 4. Female patients, with a less-dilated LV cavity, and supported with an axial-flow device were more likely to improve their myocardial function on LVAD support. To validate the identified variables, we performed a univariate linear regression analysis with each as an independent variable. LVEF, LVEDD, and HF symptoms duration (Figure 6A) appeared as significant clinical variables across both linear regression and random forest.

**Figure 6:**
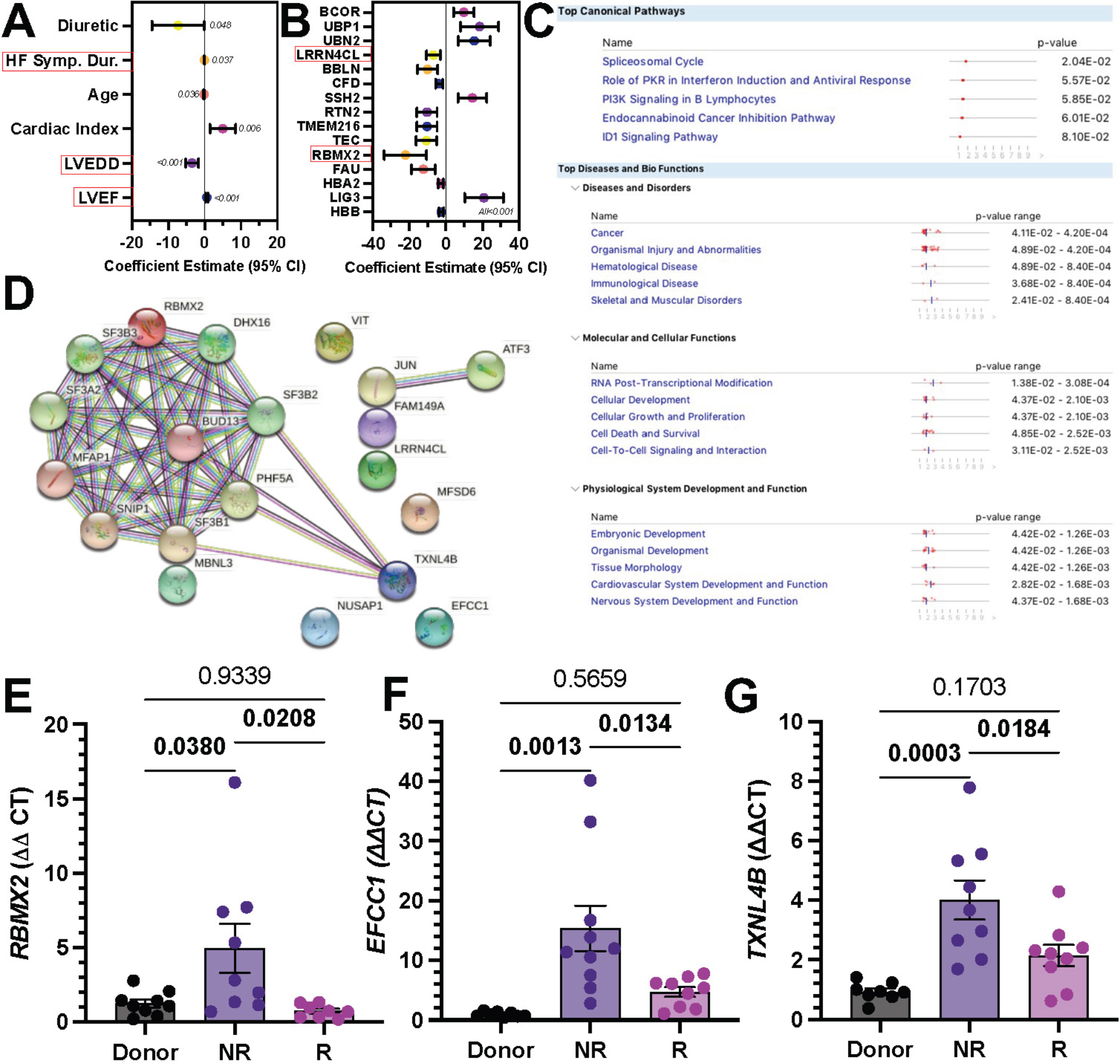
A-B) Coefficient estimates with 95% confidence intervals for the significantly identified clinical variables and mRNA transcripts associated with improved LVEF using linear regressions. Red boxes indicate variables that were identified via ML. **C)** Ingenuity Pathway Analysis shows the top canonical pathways, diseases, and bio-functions associated with the transcripts related to improved LVEF. **D)** STRING protein-protein interactions show how the proteins interact with one another. The network has been enriched with 10 additional proteins to representatively show other proteins that may be involved in improved cardiac function. **E-G)** Relative gene expression of *RBMX2*, *EFCC1* and *TXNL4B* as determined by RT-qPCR as predictive markers for improved cardiac function. Donor (non-failing human hearts), non-responders (NR), responders (R). Data is presented at mean±SEM.

**Table 3:**
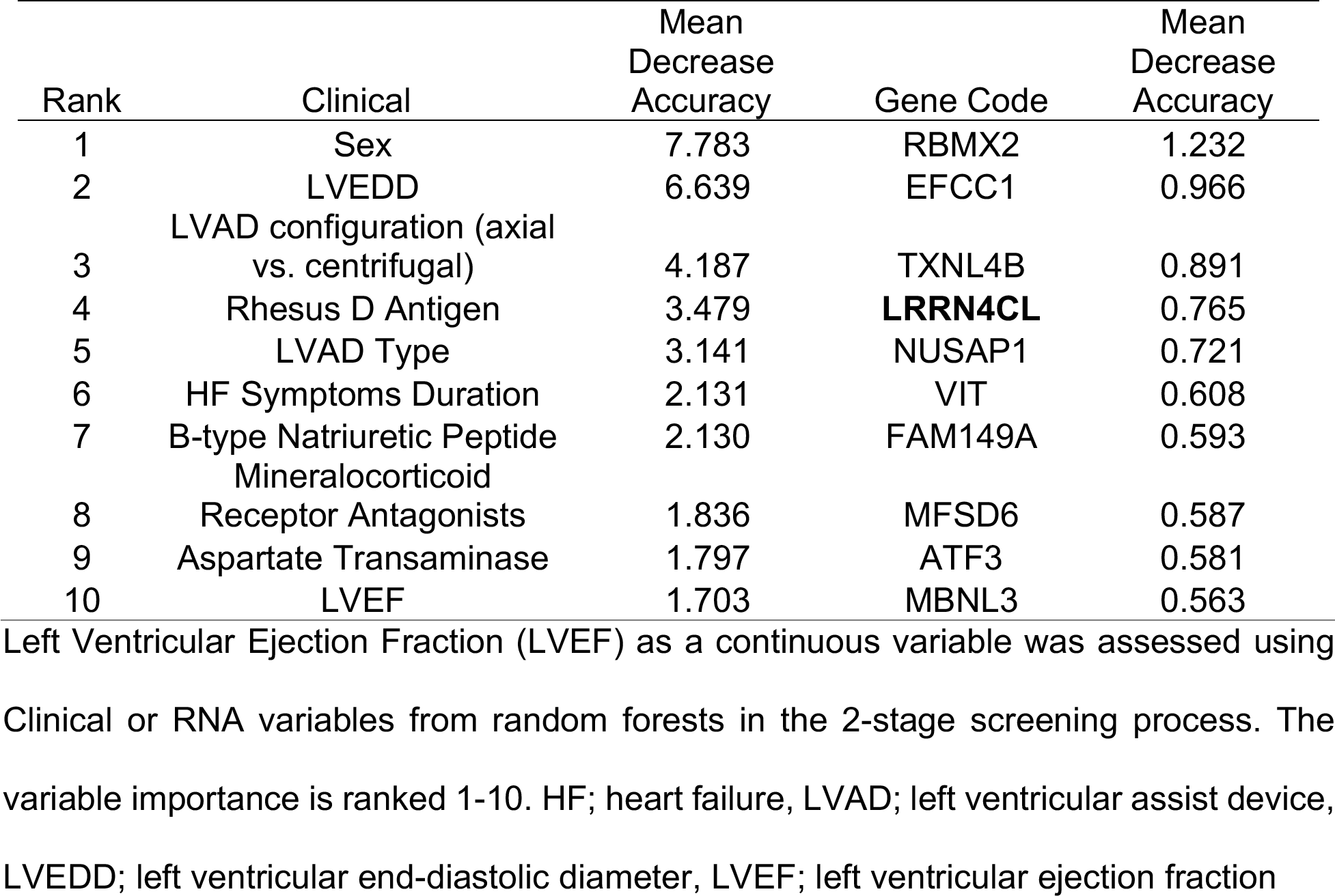
Variable importance for predicting functional myocardial improvement (increased LVEF).

The ML top-ranking RNA transcripts coded proteins such as RNA-binding motif X linked-2 (*RBMX2*), EF-Hand and Coiled-Coil Domain Containing 1 (*EFCC1*), Thioredoxin Like 4B (*TXNL4B*), and *LRRN4CL*. These specific mRNA transcripts showed a negative and non-linear relationship with the marginal prediction of improved LVEF post-LVAD. Using linear regression all of these genes showed significance (p<0.05), however only RBMX2 and LRRN4CL emerged in the top 15 with a p<0.001 (Figure 6B). Next, the ML predictive transcripts for LVEF were analyzed via IPA and STRING bioinformatics software (Figure 6C-D, respectively). IPA analysis showed the spliceosomal cycle (p=0.02) as the top canonical pathway associated with the LVEF predictive transcripts. The STRING protein-protein interactions showed nodes situated around *RBMX2* and *TXNL4B* which have been enriched with other potential interactors such as several splicing factors (*SF3A2*, *SF3B1*, *SF3B2*, *SF3B3*) and *MFAP1*.

In concordance with RNAseq analysis, RT-qPCR gene expression analysis showed *RBMX2*, *EFCC1* and *TXNL4B* had higher expression in cardiac tissue of NR compared to non-failing donors and R, which was consistent with the prediction of LVEF improvement (Figure 6E-G).

### Predicting Structural Cardiac Improvement

Cross-validation showed that the clinical data had the highest performance in predicting improvement to LVEDD using a rf model with 6-variables achieving a MAE of 0.79±0.13 on average (Figure 7). Using both data sources the 20-variable rf model attained an MAE of 0.82±0.14 on average, while using transcriptomics alone the 60 variable rf model attained an MAE of 0.96±0.19 on average.

**Figure 7:**
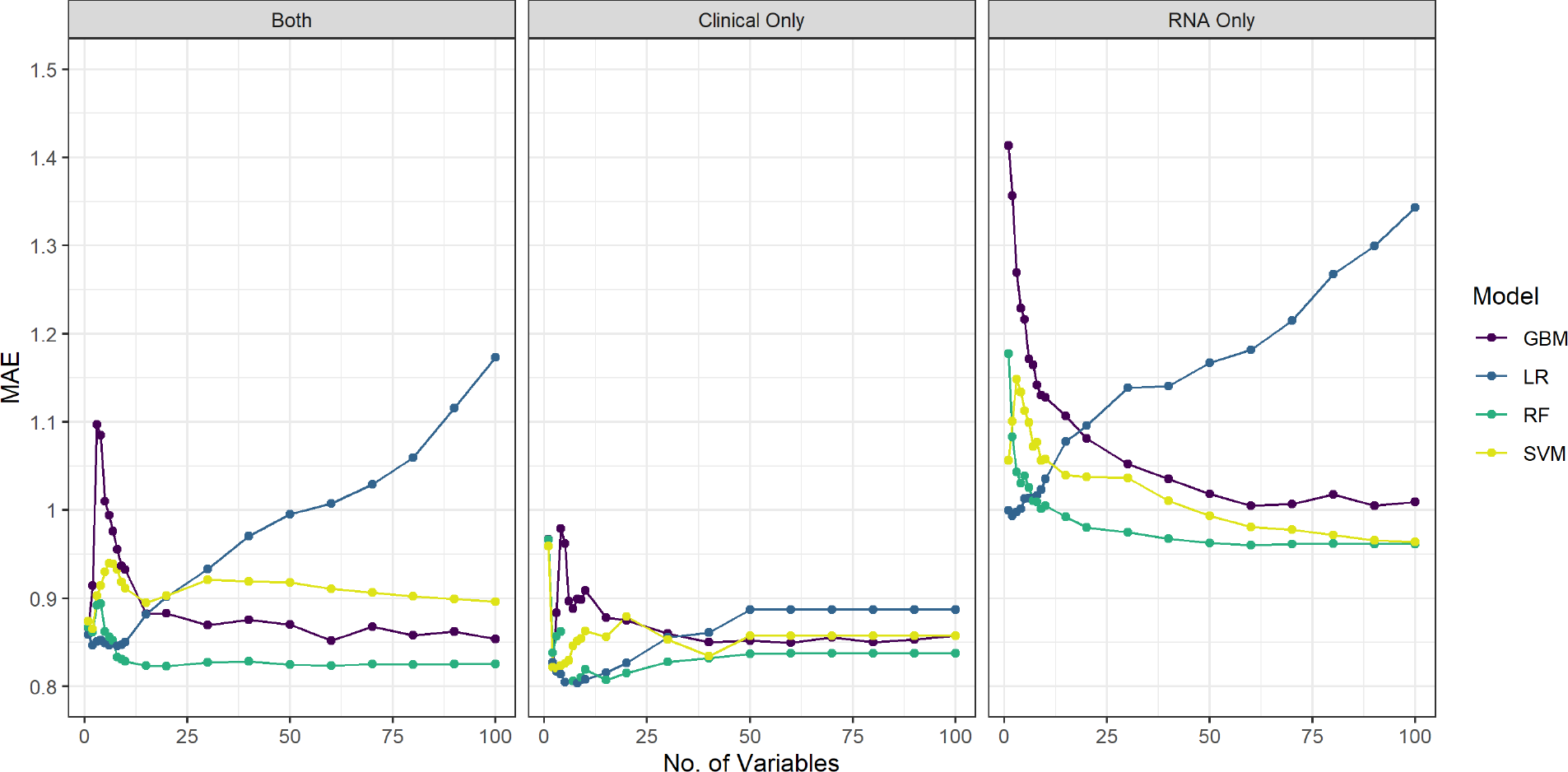
Cross-validated Mean Absolute Error (MAE) by both data sources (left), clinical characteristics (middle) or mRNA transcripts (right) and number of variables used in the model for predicting improved myocardial structure (LVEDD). A lower MAE indicates better predictive performance. Variables were selected using a two-stage screening process. Gradient Boosted Regression (gbm: 5000 trees), Logistic Regression (glm), Random Forest (rf: 5000 trees), and Support Vector Machine (svm: polynomial regression).

Baseline LVEDD, post-LVAD LVEDD, and the predicted post-LVAD LVEDD values for each patient in the testing dataset show that our predictions were conservative in predicting LVEDD, i.e., underestimated the decrease in LVEDD unless the true change from baseline was small (Left; Supplemental Figure 5). Similar to the model for LVEF, we tend to have more prediction error as the post-LVAD LVEDD moves away from the average (Right; Supplemental Figure 5).

From ML, we found that patient sex (female), baseline LVEDD, and LVAD type were the most important clinical variables for predicting LVEDD post-LVAD implant (Table 4). The partial dependency plots show that female patients, with a lower baseline LVEDD, and supported with an LVAD other than HeartWare were more likely to exhibit structural myocardial improvement on LVAD support (Supplemental Figure 6). Once more, to authenticate the identified ML variables, we performed univariate linear regression analysis which showed LVEDD, sex, ACEi/ARBs, creatinine, and platelets as significant clinical variables (Figure 8A).

**Figure 8:**
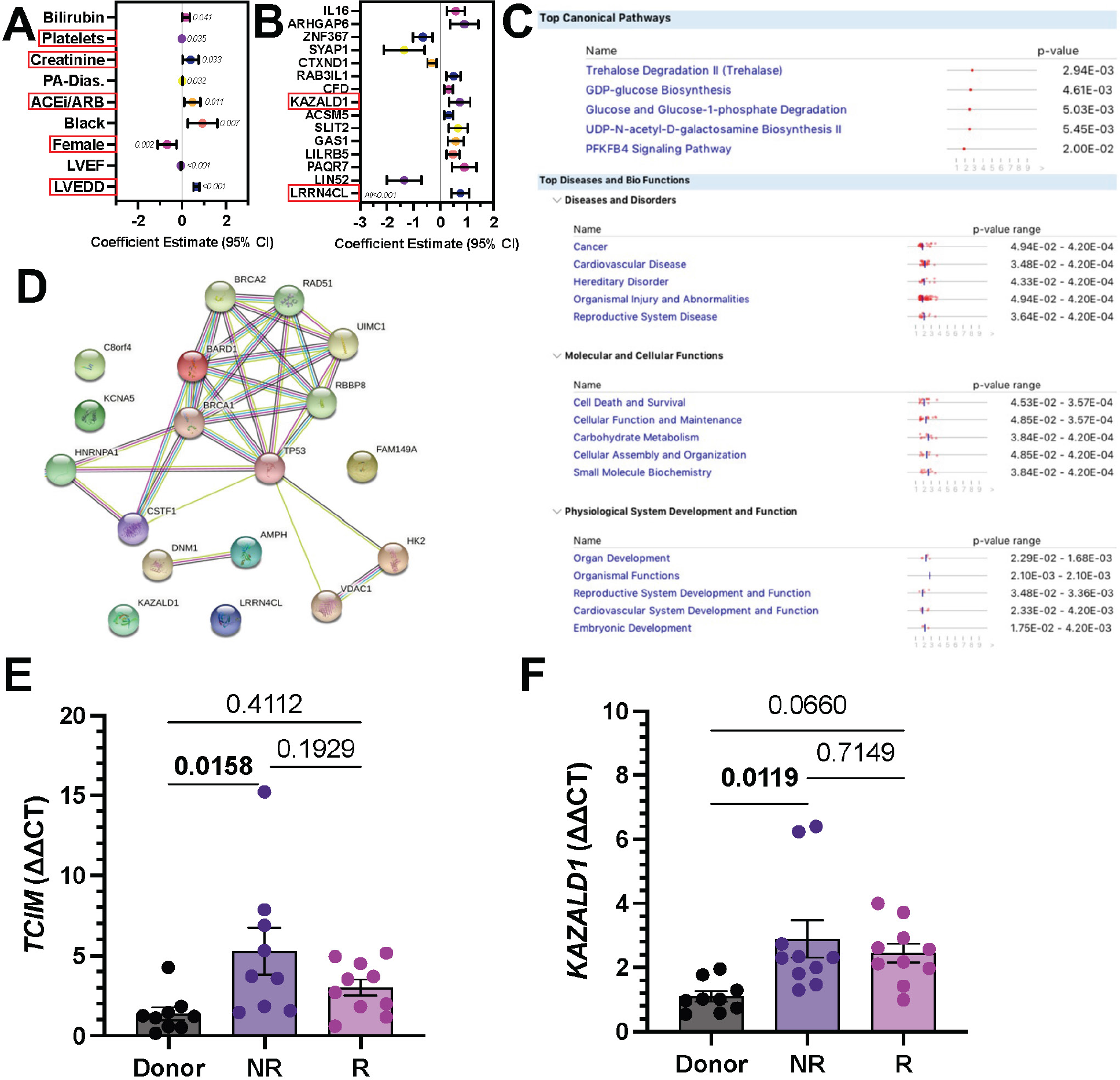
A-B) Coefficient estimates with 95% confidence intervals for the significantly identified clinical variables and mRNA transcripts associated with improved LVEDD using linear regressions. Red boxes indicate variables that were identified via ML. **C)** Ingenuity Pathway Analysis shows the top canonical pathways, diseases, and bio-functions associated with the transcripts related to improved LVEDD. **D)** STRING protein-protein interactions. The network has been enriched with 10 additional proteins to representatively show other proteins that may be involved in improved cardiac structure. **E-F)** Relative gene expression of *TCIM* and *KAZALD1* as determined by RT-qPCR as a predictive marker for improved cardiac structure. Donor (non-failing human hearts), non- responders (NR), responders (R). Data is presented at mean±SEM.

**Table 4:** Variable importance for predicting structural myocardial improvement (reduced LVEDD).

| Rank | Clinical | Mean Decrease Accuracy | Protein Name from RNA Transcript | Mean Decrease Accuracy |
| --- | --- | --- | --- | --- |
| 1 | Sex | 0.113 | TCIM | 0.014 |
| 2 | LVEDD | 0.035 | KAZALD1 | 0.010 |
| 3 | LVAD Type | 0.031 | AMPH | 0.008 |
| 4 | Platelet count | 0.022 | KCNA5 | 0.007 |
| 5 | B-type Natriuretic Peptide | 0.021 | HK2 | 0.006 |
| 6 | Rhesus D antigen | 0.017 | XIST | 0.006 |
| 7 | ACEi/ARB | 0.016 | PRKY | 0.005 |
| 8 | Creatinine | 0.015 | BARD1 | 0.004 |
| 9 | LVAD configuration | 0.011 | **LRRN4CL** | 0.004 |
| 10 | Diabetes Mellitus | 0.010 | FAM149A | 0.003 |

Left Ventricular End-Diastolic Diameter (LVEDD) was assessed as a continuous variable, using Clinical or RNA variables from random forests in the 2-stage screening process. The variable importance is ranked 1-10. ACEi; angiotensin-converting enzyme inhibitors, ARB; angiotensin receptor blocker, LVAD; left ventricular assist device, LVEDD; left ventricular end-diastolic diameter

The top-ranking ML-mRNA transcripts for predicting LVEDD improvement coded proteins such as the Transcriptional and Immune Response Regulator (*TCIM*), Kazal Type Serine Peptidase Inhibitor Domain 1 (*KAZALD1*), and again *LRRN4CL*. Our linear regression confirmed that *KAZALD1* and *LRRN4CL* were associated with improved myocardial structure (Figure 8B). IPA analysis of the ML-predictive LVEDD transcripts identified the Trehalose Degradation II (Trehalose) pathway (p=0.002, Figure 8C) as the top canonical pathway associated with improved LVEDD. The STRING protein-protein interactions showed nodes situated around *BARD1* and *HK2* (Figure 8D) which have been enriched with other potential interactors such as *BRCA1*, *VDAC1*, and *TP53*.

*TCIM* and *KAZALD1* gene expression, as assessed by RT-qPCR, was increased in failing hearts from NR and to a lesser extent from R in comparison to non-failing hearts from donors (Figure 8E). The result further validated that increased *TCIM* and *KAZALD1* expression correlated with increase in myocardial remodeling.

## DISCUSSION

Patients undergoing mechanical unloading with LVADs can experience structural and functional myocardial improvement to varying degrees, with a subset of patients exhibiting such a pronounced improvement where LVAD therapy withdrawal can be considered (4–9, 15). This clinical observation combined with the access to myocardial tissue from the LV apex at the time of LVAD implantation, provide the opportunity to perform in-depth characterizations of myocardial biology in advanced HF patients undergoing mechanical unloading (15, 32, 33). In this investigation, we used a multi-centered, and multi-model bioinformatics approach to collect human cardiac tissue at the time of LVAD implantation and characterize reverse myocardial remodeling. Specifically, we used ML (21) supplemented with logistic and linear regression analysis, to identify transcriptomics and clinical characteristics associated with LVAD-mediated myocardial recovery. To our knowledge this is the first study to combine transcriptomic and clinical data to develop predictors of myocardial recovery. LVAD-mediated mechanical unloading targeting significant cardiac reverse remodeling and subsequent device removal might be a plausible therapeutic strategy in select advanced HF patients. Our findings not only could guide the identification of advanced HF patients that could benefit from an upfront bridge-to-recovery LVAD therapeutic strategy as opposed to alternative treatment options, but importantly could help unveil the complex pathophysiological mechanisms driving myocardial recovery with implications for the broader HF population.

Patients supported with an LVAD almost universally exhibit structural cardiac improvement, but a lower proportion experiences improvement in the function of the heart (6, 8). While both criteria should be met for an LVAD-supported patient to achieve myocardial recovery and be a candidate for LVAD weaning evaluation, the dissociation between the two phenomena has not been investigated. Although it is crucial, clinically, to meet criteria for myocardial recovery based on existing cardiac functional (LVEF) and structural (LVEDD) parameter cutoffs, myocardial recovery as a biological phenomenon is not binary and, as such, we also assessed functional and structural cardiac improvement independently, as continuous variables.

Overall, we found that by using both clinical and biological data the 80-variable gbm model had the highest performance in discriminating between R and NR (binary outcome) using six and ten clinical variables had the highest performance for predicting post-LVAD LVEDD and LVEF, respectively. Although we used the unbiased rf approach as outlined by Strobl, et. al. when fitting our rf prediction models, the inferiority of the model using both data sources is likely due to some combination of overfitting and bias when splitting the variables in each decision tree (34).

### Addressing Imbalanced Responder vs. Non-Responder Class Data

In the binary R vs. NR prediction, we address the assessment of models trained and tested on imbalanced outcomes by using the PRAUC which more fairly assesses predictive performance when there is a rare outcome. We chose not to utilize any methods for correcting class imbalance, such as oversampling, undersampling, and smote since these methods have shown to negatively impact calibration and do not necessarily improve performance metrics such as AUC (35).

### Clinical Characteristics and Transcriptomics Associated with LVAD-mediated Myocardial Recovery

Our bioinformatics approach reproduced several clinical characteristics associated with LVAD-mediated myocardial recovery from previous research using traditional statistics (6, 8, 9, 16). These characteristics include duration of HF symptoms, degree of baseline LV dilation, creatinine levels, HF medical therapy prior to LVAD implantation, and support with an axial-flow LVAD (6, 8, 9, 16). The top clinical factors that were found to be discriminative of R vs NR were a shorter duration of HF symptoms, absence of medical therapy with an MRA, support with an axial-flow device or an LVAD other than HeartWare, and a higher LVEF prior to LVAD support. Shorter HF symptoms and higher LVEF may suggest a less-advanced disease stage with a higher potential for recovery. The absence of treatment with an MRA before LVAD implantation may indicate that these patients had not received the anti-remodeling effect of optimal neurohormonal blockade pre-LVAD, as has been also suggested previously (6, 9). Axial-flow devices exert a differential pattern of unloading and wall stress reduction compared to centrifugal-flow devices and have been previously associated with myocardial recovery (6, 9, 16). In our study, most patients supported with a centrifugal-flow device received a HeartWare LVAD, thus the lower chances for recovery observed with this type of LVAD might reflect a comparison with axial-flow devices. Another explanation is that the HeartWare device is considered by many as a device that could be implanted with a less invasive and shorter surgical operation. This may have led to implantation to a more critically ill population with more advanced disease and a lower recovery potential.

When we assessed the performance of the previously published ICAR score in our dataset, which predicts LVAD-mediated myocardial recovery defined as significant cardiac improvement and subsequent LVAD removal, we observed that both our 80 variable gbm model employing both clinical and transcriptomic data and our 20 variable rf model employing only clinical data, performed comparably as evidenced by the comparison of the average AUCs (p=0.80 and p=0.49, respectively). However, similar to the parsimonious ICAR score, the 20 variable rf model employing only clinical data had a similar discriminatory performance and had better calibration metrics, i.e., the predicted probability of being a R was closer to the actual observed value of R.

Alongside our clinical findings, using heart tissue before unloading, we identified a novel transcript associated with LVAD-mediated myocardial recovery. Leucine rich repeat neuronal 4CL (*LRRN4CL*) emerged as predicting LVAD-mediated myocardial recovery, improved LVEF, and improved LVEDD. From our data, *LRRN4CL* is mainly expressed in fibroblasts, and its downregulation is associated with an increased probability for LVAD-mediated myocardial recovery, improved LVEF, and improved LVEDD (Supplemental Figures 1, 2, 4, and 6, respectively). This finding was confirmed using RT-qPCR, which showed R and donor hearts had greater than a 100-fold reduced level of *LRRN4CL* compared to NR (Figure 4E). On the protein level, the abundance of LRRN4CL was lower in R when compared to NR in failing heart at the pre-LVAD time point (Supplemental Figure 2C). Additionally, our immunofluorescent staining using human cardiac tissue showed an increased signal for LRRN4CL in the failing heart of NR compared to failing heart of the R and non-failing heart of donor (Figure 4F). This expression was seen in the extracellular space between cardiomyocytes, and not intracellular. Leucine rich repeats have been reported to play a role in cardiac remodeling leading to cardiomyopathies (36, 37), however the specific role of *LRRN4CL* is undetermined. Its overexpression in NR pre-LVAD, suggest that dysregulation of *LRRN4CL* may contribute biologically to advanced HF and recovery. Future research is needed to understand the cause and effect of the changes in *LRRN4CL* expression, as it may represent an important biological target to improve both the function and structure of the heart when pharmaceutically targeted for inhibition.

### Clinical Characteristics and Transcriptomics Associated with Cardiac Functional Improvement

The top clinical variables that associated with improved cardiac function (indicated by a higher post-LVAD LVEF as a continuous variable) were female sex, lower baseline LVEDD, support with an axial-flow device or an LVAD other than HeartWare, and negative Rhesus D antigen status. LV chamber size may indicate the extent of myocardial remodeling and reflect the chronicity of disease, as patients with a less dilated LV have been previously identified as having a higher propensity for myocardial recovery (6, 8, 9, 16). Similarly, female sex has been previously associated with an increased likelihood of reverse myocardial remodeling in chronic systolic HF (38).

Transcriptomics and RT-qPCR revealed that low expression of *RBMX2 (*RNA binding motif protein x-linked 2)*, EFCC1* (EF-hand and coiled-coil domain containing 1) and *TXNL4B* (thioredoxin like 4B) differentiated NR from R and predicted improved cardiac function following unloading (Figure 6E-G). *RBMX2* is a gene involved in pre-mRNA splicing as a member of the activated spliceosome, but little is known about its role in HF. *EFCC1* is a member of the coiled-coil domain containing proteins with a diverse array of functions including calcium ion binding (39, 40). *EFCC1* has been implicated in several pathological diseases including mitochondrial disease, cancer, and diabetes (39). *TXNL4B*, also known as *DIM2*, has been reported to be involved in pre-mRNA splicing, an important physiological mechanism which generates transcriptional diversity, regulates gene expression (41, 42), and is required for cell cycle progression (43). At the same time, it has been reported that mRNA splicing is altered in human heart disease with a mis-regulation and decreased efficiency for splicing (44). *RBMX2*, *EFCC1*, and *TXNL4B* transcripts were differentially expressed between R and NR pre-LVAD and their signature suggested that these genes may contribute to regulation of myocardial contractility.

### Clinical Characteristics and Transcriptomics Associated with Cardiac Structural Improvement

We and others have previously demonstrated that structural myocardial improvement takes place in a much higher proportion of LVAD patients (compared to cardiac functional improvement) (5, 6, 8). Therefore, we sought to explore whether the transcriptomics and clinical characteristics associated with structural improvement will be different than those associated with functional improvement. Female sex, lower baseline LVEDD, support with an LVAD other than HeartWare, as well as higher platelet count and B-type natriuretic peptide levels were the top clinical variables associated with structural myocardial improvement. An elevated platelet count prior to LVAD implantation was more common among patients experiencing LVAD-mediated myocardial recovery in prior studies (6). Higher B-type natriuretic peptide levels suggest more pronounced LV wall stress due to pressure and volume overload, the attenuation of which upon mechanical unloading might lead to a higher degree of structural reverse remodeling.

In combination with the clinical factors associated with structural reverse remodeling, our transcriptomic data revealed that lower *TCIM* (*TC1*, or *C8orf4*) expression pre-LVAD predicted improved LVEDD following LVAD unloading. *TCIM* has been shown to act as an inflammatory regulator (45, 46, 47) which enhances *NFKB* transcriptional activity, suggesting an augmentation of inflammatory signals and cellular stress (48). Our predictions indicated that an increased expression of *TCIM* in HF was associated with adverse cardiac remodeling, and a potential link between reduced inflammatory signals through lower *TCIM* expression and improved structural reverse remodeling (8). Moreover, *KAZALD1* was also identified through ML and linear regression analysis as being a predictor for improved LVEDD, and RT-qPCR showed that increase *TCIM* and *KAZALD1* associated with increase in LVEDD in failing heart from both NR and R at pre-LVAD.

## STUDY LIMITATIONS

Our sample size of 208 patients was modest which in combination with the relatively low proportion of LVAD-supported patients achieving myocardial recovery limits the stability of the feature effect estimates and leads to potential overfitting, even at smaller feature subsets. Given our small sample size, we felt it more appropriate to derive the model on a large training set in cross-validation (90%) rather than designate a hold-out set or derive on a smaller proportion of our data with the understanding that this would lead to a larger variance in the cross-validated results. However, by training on the majority of our data, the results should be most similar to if we had trained a model on our full data and tested on an external set with similar properties as our data. Additionally, given the size of the transcriptomics feature set, we utilized two screening steps, one using regression and one using ML, in order to limit computational time of fitting models and to avoid overfitting. This two-step process may have biased the models towards transcriptomic features with a linear relationship with the outcome, though exploratory analysis showed a linear relationship was typical. Lastly, we were interested in predicting recovery, and therefore did not incorporate transcriptomics data post-LVAD unloading which may lead to a more refined predictive model for myocardial recovery due to the adaptive and maladaptive biological responses that can occur following LVAD support.

## CONCLUSION

In summary, to our knowledge we are the first to employ a bioinformatics approach combining clinical variables and omics data from human HF patients undergoing LVAD implantation to create models predicting LVAD-mediated myocardial recovery. The convergence of clinical characteristics with biological information derived from the analysis of myocardial tissue, exhibited a superior predictive capacity which will only be optimized in the future as new data becomes available and our understanding of HF evolves. Additionally, we identified and confirmed novel transcripts such as *LRRN4CL* (ligand binding and programmed cell death), *RBMX2* and *TXNL4B* (both mRNA splicing), and *EFCC1* (calcium ion binding) in the failing myocardium at the time of LVAD implantation that differentiates responders from non-responder and will guide future innovative research. These results illuminate several important clinical and molecular drivers of LVAD-mediated myocardial recovery, and unveil potential mechanistic pathways that have a promising future in the diagnostic, prognostic, and therapeutic approaches for the broader population of patients suffering from HF.

## AUTHOR CONTRIBUTIONS

J.R.V., C.P.K., B.J.B., W.L.H, S.A.S, G.S.D, S.N., P.S., M.S.S., M.K., and S.G.D. designed the research studies. J.R.V., C.P.K., B.J.B., Y.H., I.T., R.B., B.H., T.S.S., J.L., R.H., E.T., S.N., and O.W.P. conducted experiments. J.R.V., C.P.K., B.J.B., Y.H., I.T., R.B., T.S.S., J.L., R.H., E.T., S.N., O.W.P., S.C.K., T.C.H., S.M., S.B., M.S., R.A., T.H.G., C.H.S., P.S., M.K., and S.G.D. acquired data. J.R.V., C.P.K., B.J.B., Y.H., I.T., R.B., T.S.S., J.L., R.H., E.T., S.N., O.W.P., S.C.K., T.C.H., S.M., S.B., M.S., R.A., T.H.G., C.H.S., P.S., M.S.S., M.K., and S.G.D. analyzed data. J.R.V., C.P.K., B.J.B., R.B., S.N., S.C.K., T.C.H., S.M., S.B., P.S., M.K., and S.G.D. wrote the manuscript. J.R.V., C.P.K., B.J.B., Y.H., I.T., R.B., T.S.S., J.L., R.H., E.T., S.N., W.L.H, S.A.S, G.S.D, O.W.P., S.C.K., T.C.H., S.M., S.B., M.S., R.A., T.H.G., C.H.S., P.S., M.S.S., M.K., and S.G.D. reviewed and approved the manuscript. J.R.V., B.J.B, and C.P.K share the first author position.

## Supporting information

Supplement_Visker 2024

## ACKNOWLEDGEMENTS

We thank Timothy Parnell for technical assistance and RNA Sequencing data analysis. We thank the Genomics core at the University of Utah for performing RNA sequencing and help with RT-qPCR. We also thank all the lab members and summer undergraduate trainees for sample collection and processing.

## FUNDING

We acknowledge the following for funding: R01HL135121 and R01HL132067 to SGD; the Nora Eccles Treadwell Foundation to S.G.D.; the AHA Heart Failure Strategically Focused Research Network 16SFRN29020000 to S.G.D. and CHS Merit Review Award I01 CX002291 U.S. Dept. of Veterans Affairs to S.G.D. This research was supported by the National Institutes of Health under Ruth L. Kirschstein National Research Service Award 5T32DK091317 to J.R.V. (National Institute of Diabetes and Digestive and Kidney Diseases), 2T32HL007576-36 to C.P.K. and 5T32HL007576-33 to I.T. (National Heart, Lung, and Blood Institute). This investigation was supported by the University of Utah Population Health Research (PHR) Foundation, with funding in part from the National Center for Advancing Translational Sciences of the National Institutes of Health under Award Number UL1TR002538. P.S. is supported by an NIH Career Development Award 1K23HL143179. The content is solely the responsibility of the authors and does not necessarily represent the official views of the National Institutes of Health.

## Notes

### Competing Interest Statement

Dr. Drakos serves as a consultant for Abbott Laboratories and has received research support from Novartis. Dr. Summers is a co-founder and consultant with Centaurus Therapeutics, Inc. The remaining authors have nothing to disclose.

## REFERENCES

1. Metra M, Carubelli V, Ravera A, Stewart Coats AJ. Heart failure 2016: still more questions than answers. Int J Cardiol. 2017 Jan 15;227:766-777. doi: 10.1016/j.ijcard.2016.10.060.

2. Molina EJ, Shah P, Kiernan MS, et. al. The Society of Thoracic Surgeons Intermacs 2020 Annual Report. Ann Thorac Surg. 2021 Mar;111(3):778-792. doi: 10.1016/j.athoracsur.2020.12.038.

3. Birks EJ, Tansley PD, Hardy J, et. al. Left ventricular assist device and drug therapy for the reversal of heart failure. N Engl J Med. 2006 Nov 2;355(18):1873–1884. doi: 10.1056/NEJMoa053063.

4. Birks EJ, George RS, Hedger M, et. al. Reversal of severe heart failure with a continuous-flow left ventricular assist device and pharmacological therapy: a prospective study. Circulation. 2011 Feb 1;123(4):381–390. doi: 10.1161/CIRCULATIONAHA.109.933960.

5. Drakos SG, Wever-Pinzon O, Selzman CH, et. al. Magnitude and time course of changes induced by continuous-flow left ventricular assist device unloading in chronic heart failure: insights into cardiac recovery. J Am Coll Cardiol. 2013 May 14;61(19):1985–1994. doi: 10.1016/j.jacc.2013.01.072.

6. Topkara VK, Garan AR, Fine B, et. al. Myocardial Recovery in Patients Receiving Contemporary Left Ventricular Assist Devices: Results From the Interagency Registry for Mechanically Assisted Circulatory Support (INTERMACS). Circ Heart Fail. 2016 Jul;9(7):10.1161/CIRCHEARTFAILURE.116.003157 e003157. doi: 10.1161/CIRCHEARTFAILURE.116.003157.

7. Birks EJ, Drakos SG, Patel SR, et. al. Prospective Multicenter Study of Myocardial Recovery Using Left Ventricular Assist Devices (RESTAGE-HF [Remission from Stage D Heart Failure]): Medium-Term and Primary End Point Results. Circulation. 2020 Nov 24;142(21):2016–2028. doi: 10.1161/CIRCULATIONAHA.120.046415.

8. Shah P, Psotka M, Taleb I, et. al. Framework to Classify Reverse Cardiac Remodeling With Mechanical Circulatory Support: The Utah-Inova Stages. Circ Heart Fail. 2021 May;14(5):e007991. doi: 10.1161/CIRCHEARTFAILURE.120.007991.

9. Wever-Pinzon O, Drakos SG, McKellar SH, et. al. Cardiac Recovery During Long-Term Left Ventricular Assist Device Support. J Am Coll Cardiol. 2016 Oct 4;68(14):1540–1543.

10. Margulies KB. Reversal mechanisms of left ventricular remodeling: Lessons from left ventricular assist device experiments. J Card Fail. 2002 Dec;8(6 Suppl):S500-505. doi: 10.1054/jcaf.2002.129264.

11. Terracciano CM, Hardy J, Birks EJ, Khaghani A, Banner NR, Yacoub MH. Clinical Recovery From End-Stage Heart Failure Using Left-Ventricular Assist Device and Pharmacological Therapy Correlates With Increased Sarcoplasmic Reticulum Calcium Content but Not With Regression of Cellular Hypertrophy. Circulation. 2004 May 18;109(19):2263–2265. doi: 10.1161/01.CIR.0000129233.51320.92.

12. Diakos NA, Navankasattusas S, Abel ED, et. al. Evidence of Glycolysis Up- Regulation and Pyruvate Mitochondrial Oxidation Mismatch During Mechanical Unloading of the Failing Human Heart: Implications for Cardiac Reloading and Conditioning. JACC Basic Transl Sci. 2016 Oct;1(6):432–444. doi: 10.1016/j.jacbts.2016.06.009.

13. Badolia R, Ramadurai DK, Abel ED, et. al. The Role of Non-Glycolytic Glucose Metabolism in Myocardial Recovery Following Mechanical Unloading and Circulatory Support in Chronic Heart Failure. Circulation. 2020 Jul 21;142(3):259-274. doi: 10.1161/CIRCULATIONAHA.119.044452.

14. Cluntun AA, Badolia R, Lettlova S, et. al. The pyruvate-lactate axis modulates cardiac hypertrophy and heart failure. Cell Metab. 2021 Mar 2;33(3):629–648.e10. doi: 10.1016/j.cmet.2020.12.003.

15. Drakos SG, Pagani FD, Lundberg MS, Baldwin JT. Advancing the Science of Myocardial Recovery With Mechanical Circulatory Support: A Working Group of the National, Heart, Lung, and Blood Institute. JACC Basic Transl Sci. 2017 Jun;2(3):335–340. doi: 10.1016/J.jacbts.2016.12.003.

16. Antonides CF, Schoenrath F, de By TM, et. al. Outcomes of patients after successful left ventricular assist device explantation: a EUROMACS study. ESC Heart Fail. 2020 Jun;7(3):1085–1094. doi: 10.1002/ehf2.12629.

17. Seidel T, Navankasattusas S, Ahmad A, et. al. Sheet-Like Remodeling of the Transverse Tubular System in Human Heart Failure Impairs Excitation-Contraction Coupling and Functional Recovery by Mechanical Unloading. Circulation. 2017 Apr 25;135(17):1632–1645. doi: 10.1161/CIRCULATIONAHA.116.024470.

18. Diakos NA, Taleb I, Kyriakopoulos CP, et. al. Circulating and Myocardial Cytokines Predict Cardiac Structural and Functional Improvement in Patients With Heart Failure Undergoing Mechanical Circulatory Support. J Am Heart Assoc. 2021 Oct 19;10(20):e020238. doi: 10.1161/JAHA.120.020238.

19. Yerushalmi R, Woods R, Ravdin PM, Hayes MM, Gelmon KA. Ki67 in breast cancer: prognostic and predictive potential. Lancet Oncol. 2010 Feb;11(2):174–183. doi: 10.1016/S1470-2045(09)70262-1.

20. Payne SJ, Bowen RL, Jones JL, Wells CA. Predictive markers in breast cancer-- the present. Histopathology. 2008 Jan;52(1):82–90. doi: 10.1111/j.1365-2559.2007.02897.x.

21. Poss AM, Maschek JA, Cox JE, et. al. Machine learning reveals serum sphingolipids as cholesterol-independent biomarkers of coronary artery disease. J Clinl Invest. 2020 Mar 2;130(3):1363–1376. doi: 10.1172/JCI131838.

22. Jurmeister P, Bockmayr M, Seegerer P, et. al. Machine learning analysis of DNA methylation profiles distinguishes primary lung squamous cell carcinomas from head and neck metastases. Sci Transl Med. 2019 Sep 11;11(509):eaaw8513. doi: 10.1126/scitranslmed.aaw8513.

23. Murray PG, Stevens A, Leonibus CD, Koledova E, Chatelain P, Clayton PE. Transcriptomics and machine learning predict diagnosis and severity of growth hormone deficiency. JCI Insight. 2018 Apr 5;3(7):e93247. doi: 10.1172/jci.insight.93247.

24. Hathaway QA, Roth SM, Pinti MV, et. al. Machine-learning to stratify diabetic patients using novel cardiac biomarkers and integrative genomics. Cardiovasc Diabetol. 2019 Jun 11;18(1):78. doi: 10.1186/s12933-019-0879-0.

25. Lang RM, Badano LP, Mor-Avi V, et. al. Recommendations for cardiac chamber quantification by echocardiography in adults: an update from the American Society of Echocardiography and the European Association of Cardiovascular Imaging. Eur Heart J Cardiovasc Imaging. 2015 Mar;16(3):233–270. doi: 10.1093/ehjci/jev014.

26. Azur MJ, Stuart EA, Frangakis C, Leaf PJ. Multiple imputation by chained equations: what is it and how does it work? Int J Methods Psychiatr Res. 2011 Mar;20(1):40–49. doi: 10.1002/mpr.329.

27. Wood AM, White IR, Royston P. How should variable selection be performed with multiply imputed data? Stat Med. 2008 Jul 30;27(17):3227–3246. doi: 10.1002/sim.3177.

28. Van Calster B, McLernon DJ, Van Smeden M, et. al. Calibration: the Achilles heel of predictive analytics. BMC Med. 2019 Dec 16;17(1):230. doi: 10.1186/s12916-019-1466-7.

29. Cai W, Liu T, Xue X, et. al. CT Quantification and Machine-learning Models for Assessment of Disease Severity and Prognosis of COVID-19 Patients. Acad Radiol. 2020 Dec;27(12):1665-1678. doi: 10.1016/j.acra.2020.09.004.

30. R Development Core Team. R: A Language and Environment for Statistical Computing. *R Foundation for Statistical Computing*, Vienna, Austria, 2014.

31. Amrute JM, Luo X, Penna V, et. al. Targeting the Immune-Fibrosis Axis in Myocardial Infarction and Heart Failure. BioRxiv. 2022.10.17.512589.

32. Taleb I, Tseliou E, Fang JC, et. al. A Mechanical Bridge to Recovery as a Bridge to Discovery: Learning From Few and Applying to Many. Circulation. 2022 Feb 22; 145(8):562–564. doi:10.1161/CIRCULATIONAHA.120.052141.

33. Kyriakopoulos CP, Kapelios CJ, Stauder EL, et. al. LVAD as a Bridge to Remission from Advanced Heart Failure: Current Data and Opportunities for Improvement. J Clin Med. Jun 20;11(12):3542. doi: 10.3390/jcm11123542.

34. Strobl C, Boulesteix AL, Zeileis A, Hothorn T. Bias in random forest variable importance measures: illustrations, sources and a solution. BMC Bioinformatics. 2007 Jan 25;8:25. doi: 10.1186/1471-2105-8-25.

35. van den Goorbergh, van Smeden M, Timmerman D, Van Calster B. The harm of class imbalance corrections for risk prediction models: illustration and simulation using logistic regression. J Am Med Inform Assoc. 2022. Aug 16;29(19):1525- 1534.

36. Li R, Fang J, Huo B, et. al. Leucine-rich repeat neuronal protein 4 (LRRN4) potentially functions in dilated cardiomyopathy. Int J Clin Exp Pathol. 2017 Sep 1;10(9):9925-9933. eCollection 2017.

37. Brody MJ, Lee Y. The Rold of Leucine-Rich Repeat Containing Protein 10 (LRRC10) in Dilated Cardiomyopathy. Front Physiol. 2016 Aug 3;7:337. doi: 10.3389/fphys.2016.00337.

38. Aimo A, Vergaro G, Castiglione V, et. al. Effect of Sex on Reverse Remodeling in Chronic Systolic Heart Failure. JACC. Heart Fail. 2017 Oct;5:735–742. doi: 10.1016/j.jchf.2017.07.011.

39. Xia L, Zhu Y, Zhang C, et. al. Decreased expression of EFCC1 and its prognostic value in lung adenocarcinoma. Ann Transl Med. Nov;7(22):672. doi: 10.21037/atm.2019.10.41.

40. Burkhard P, Stetefeld J, Strelkov SV. Coiled coils: a highly versatile protein folding motif. Trends Cell Biol. 2001 Feb; 11(2):82–88. doi:10.1016/s0962-8924(00)01898-5.

41. Maatz H, Jens M, Liss M, et. al. RNA-binding protein RBM20 represses splicing to orchestrate cardiac pre-mRNA processing. J Clin Invest. 2014 Aug;124:3419- 3430. doi: 10.1172/JCI74523.

42. Davis FJ, Gupta M, Pogwizd SM, et. al. Increased expression of alternatively spliced dominant-negative isoform of SRF in human failing hearts. Am J Physiol Heart Circ Physiol. 2002 Apr;282(4):H1521–1533. doi: 10.1152/ajpheart.00844.2001.

43. Sun X, Zhang H, Wang D, et. al. DLP, a novel Dim1 family protein implicated in pre-mRNA splicing and cell cycle progression. J Biol Chem. 2004 Jul 30; 279(31):32839-32847. doi: 10.1074/jbc.M402522200.

44. Kong SW, Hu YW, Ho JW, et. al. Heart failure-associated changes in RNA splicing of sarcomere genes. Circ Cardiovasc Genet. 2010 Apr; 3(2):138–146. doi: 10.1161/CIRCGENETICS.109.904698.

45. Sunde M, McGrath KC, Young L, et. al. TC-1 is a novel tumorigenic and natively disordered protein associated with thyroid cancer. Cancer Res. 2004 Apr 15;64(8):2766-2773. doi: 10.1158/0008-5472.can-03-2093.

46. Kim B, Koo H, Yang S, et. al. TC1(C8orf4) correlates with Wnt/beta-catenin target genes and aggressive biological behavior in gastric cancer. Clin Cancer Res. 2006 Jun 1;12 (11 Pt 1):3541-3548. doi: 10.1158/1078-0432.CCR-05-2440.

47. Park J, Jung Y, Kim J, et. al. TC1 (C8orf4) is upregulated by cellular stress and mediates heat shock response. Biochem Biophys Res Commun. 2007 Aug 24;360(2):447-452.

48. Kim J, Kim Y, Kim HT, et. al. TC1(C8orf4) is a novel endothelial inflammatory regulator enhancing NF-kappaB activity. J Immunol. 2009 Sep 15;183(6):3996- 4002. doi: 10.4049/jimmunol.0900956.

