## Supplement_Visker 2024 for "Integrating molecular and clinical variables to predict myocardial recovery"

### SUPPLEMENTAL MATERIALS

#### RNA Isolation

RNA yield and purity (A260/A280 > 2.0) were determined by spectrophotometric absorbance (NanoDrop™ Lite Spectrophotometer, Thermo Scientific™: Waltham, MA, USA). The RNA-sequencing library was prepared using the Illumina TruSeq Stranded RNA kit with Ribo-Zero Gold to remove rRNA and sequenced on an Illumina HiSeq 2500 with 50 bp single-end reads. RNA sequence reads were aligned to the human genome version hg38 with Novoalign (version 2.8.1) against an index containing genome sequences plus all splice junctions (known and theoretical, 46 bp radius) generated with USeq MakeTranscriptome (version 8.8.1) and Ensembl gene annotation (release 84), allowing for up to 50 alignments per read. Raw alignments were converted to genomic coordinates using USeq 9. SamTranscriptomeParser (version 8.8.8), allowing for a maximum of 1 alignment per read. RNA-Seq quality was evaluated with results from Picard CollectRnaSeqMetrics (version 1.137). Gene counts were collected using Sub Read feature Counts (version 1.5.1) and Ensembl GRCh38 annotation (release 87) in a stranded fashion, assigning to genes with the largest overlap. Low and non-expressed transcripts were removed when the maximum observed count across all samples was ≤ 10 counts. Differentially expressed transcripts were identified using DESeq2 (version 1.16.1).

#### Quantitative Real Time Polymerase Chain Reactions (RT-qPCR)

Complimentary deoxyribonucleic acid (cDNA) was synthesized using oligo (dT) primers with the Luna® Universal qPCR Master Mix cDNA Synthesis Kit (New England

Biolabs, Ipswich, MA). Relative gene expression differences for *LRRN4CL* (Assay ID: Hs04972223_s1), *EFCC1* (Assay ID: Hs00227049_m1), *TXNL4B* (Assay ID: Hs04188484_m1), and *TCIM* (Assay ID: Hs00535539_s1) were detected by using the QuantStudio 12k Flex Real-Time PCR System (Applied Biosystems, Thermo Fisher Scientific) using Thermo Fisher Taqman® gene expression assays. The results were normalized against *Vinculin* (Assay ID: Hs00419715_m1) gene expression using the ΔΔCT method.

**Western Blotting**

After determining total protein concentration using the BCA assay, samples were heated for 5 minutes at 95°C. 30-50 μg of protein lysate were resolved on SDS polyacrylamide gel according to the standard procedure of 20 mA per gel and blotted onto a nitrocellulose membrane 0.45 μm (GE Healthcare) via Mini Trans-blot module (Bio-Rad) at a constant voltage of 100 V for 2 hours. Next, membranes were blocked with 5% proteomic grade non-fat dried milk (NFDM) for 1 hour and then the membranes were incubated overnight in 5% bovine serum albumin. Antibodies probing for LRRN4CL (abcam 1:500) with a loading control of Vinculin (cell signaling 1:1000) were used. Following overnight incubation, membranes were washed with TBS-T and placed in an incubation with a fluorophore conjugated secondary antibody (Rockland Immunochemical, 1:10,000) in 1% NFDM with TBS-T for 1 hour. Next, membranes were washed again with TBS-T and fluorescence was detected using an Odyssey CLx imaging system (LI-COR Biosciences). Values are represented as arbitrary units (au) by normalizing LRRN4CL to Vinculin.

**Immunofluorescent Confocal Microscopy**

Human heart tissues were used for this study. Cardiac tissues were fixed using 4% paraformaldehyde (PFA) followed by paraffin embedding. Tissues were sectioned at 8μm thickness, placed on a glass coverslip, deparaffinized, and heat-induced epitope retrieval (HEIR) was performed. Following HEIR, tissue staining was completed using wheat germ agglutin-conjugated to Alexa 555 (WGA, 1:1000), and LRRN4CL (abcam 188018, 1:100). The secondary antibody of chicken-anti-rabbit Alexa 488 was used. Then, sections were placed in a mounting medium supplemented with DAPI and stored at 4° Celsius until imaging on the following day. All images were acquired using a Leica DMi8 (Leica Microsystems, Germany) confocal laser microscope with the same laser settings across all samples and processed using Leica Application Suite X.

**Predictive Models and Assessing Performance**

In the traditional screening stage, for the LVAD-responder vs. non-responder groups, a separate logistic regression was performed using each candidate predictor, and for the LVEF and LVEDD outcomes, a simple linear regression was performed using each candidate predictor. The candidate variables were then ranked in ascending order according to p-values, and the top 2.5% of candidate variables brought forth to the next screening stage. For the clinical variables, all clinical variables were considered and brought to the second stage. In the ML screening stage, the importance of each of the top 2.5% of variables brought forward from the first screening stage was computed using the random forest conditional permutation algorithm (47) (with the square root of total candidates randomly selected for each split, 5000 trees). We considered the following types of models for prediction: logistic regression (glm), random forest (rf, 5000 trees), gradient boosted regression trees (gbm, 5000 trees), and support vector machine (svm, polynomial regression) using the top 5, 10, 15, 20, 40, 60, 80, or 100 variables from the $2^{nd}$ stage of screening.

### SUPPLEMENTAL FIGURES AND TABLES

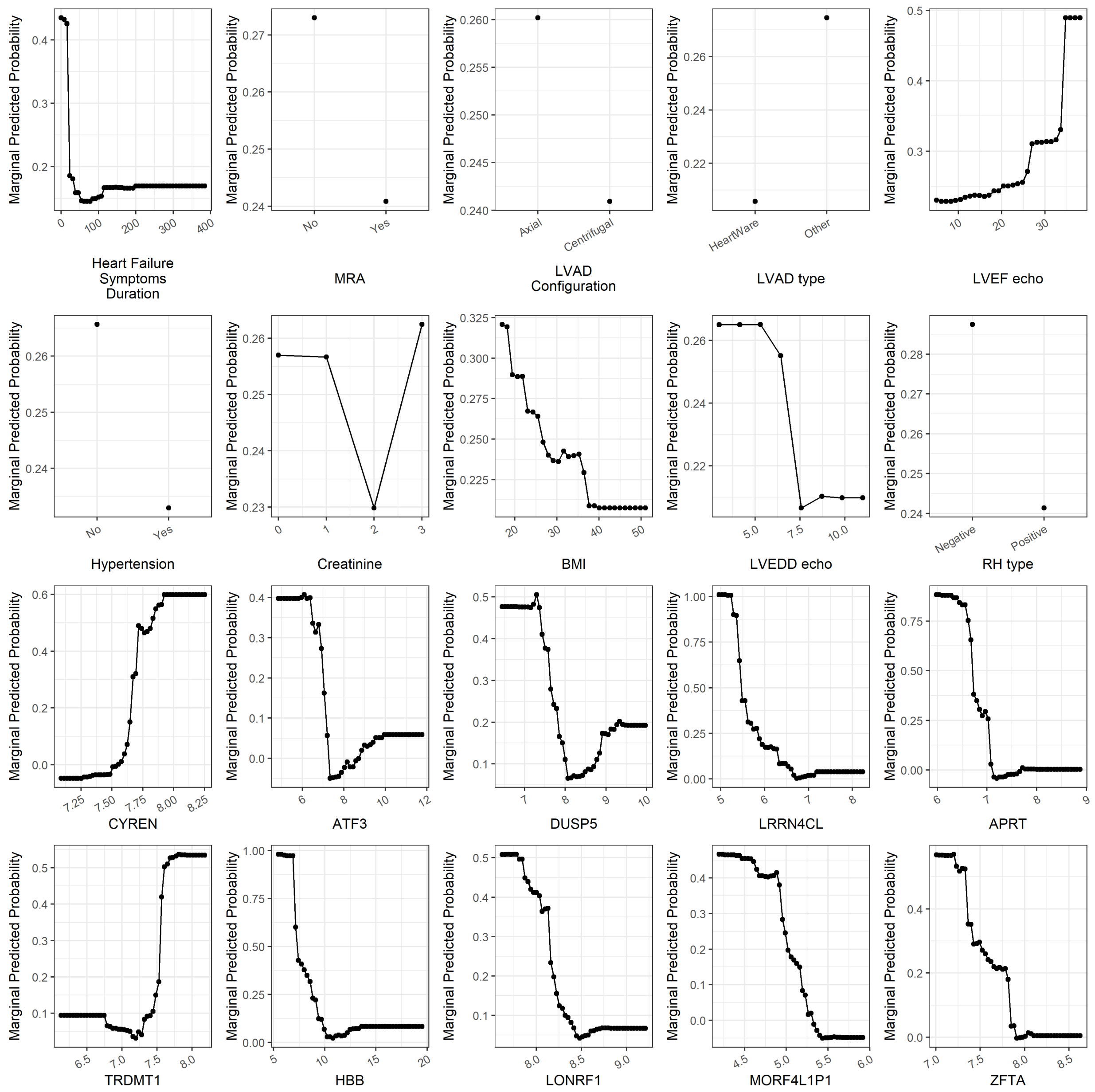


**Supplemental Figure 1**: Partial Dependency Plots for the prediction of LVAD-mediated myocardial recovery (LVAD-responder). From the Random Forest (rf) two-stage screening process, the top 10 clinical and mRNA-transcript variables are shown. Y-Axis: An increased marginal predicted value indicates a greater probability for a variable to predict responders. X-Axis: Heart Failure Symptoms Duration (Months), MRA (No vs. Yes), LVAD Configuration (Axial vs. Centrifugal), LVAD type (HeartWare vs. Other), LVEF echo (%), Hypertension (No vs. Yes), Creatinine (mg/dL), BMI (kg/m$^2$), LVEDD echo (cm), RH type (Negative vs. Positive), *CYREN*, *ATF3*, *DUSP5*, *LRRN4CL*, *APRT*, *TRDMT1*, *HBB*, *LONRF1*, *MORF4L1P1*, *ZFTA* (gene expression represented by R-Log Values).

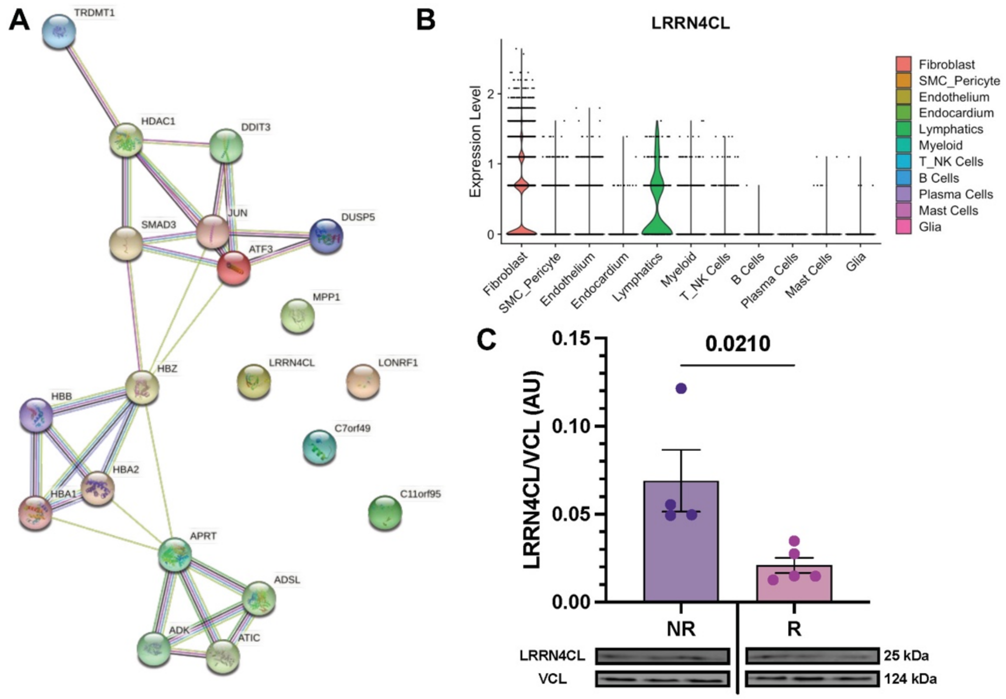


**Supplemental Figure 2: A)** STRING protein-protein interactions show how the proteins interact with one another. The network has been enriched with 10 additional proteins to representatively show other proteins that may be involved in LVAD-mediated myocardial recovery. **B)** Differential cellular expression of LRRN4CL from cellular indexing of transcriptomes and epitomes by sequencing (CITE-seq) in 22 explanted human hearts from healthy donors, acute myocardial infarction, and chronic ischemic and non-ischemic cardiomyopathy patients. **C)** Relative protein abundance, determined via western blotting, of LRRN4CL normalized to Vinculin in non-responders and responders pre-LVAD implantation.

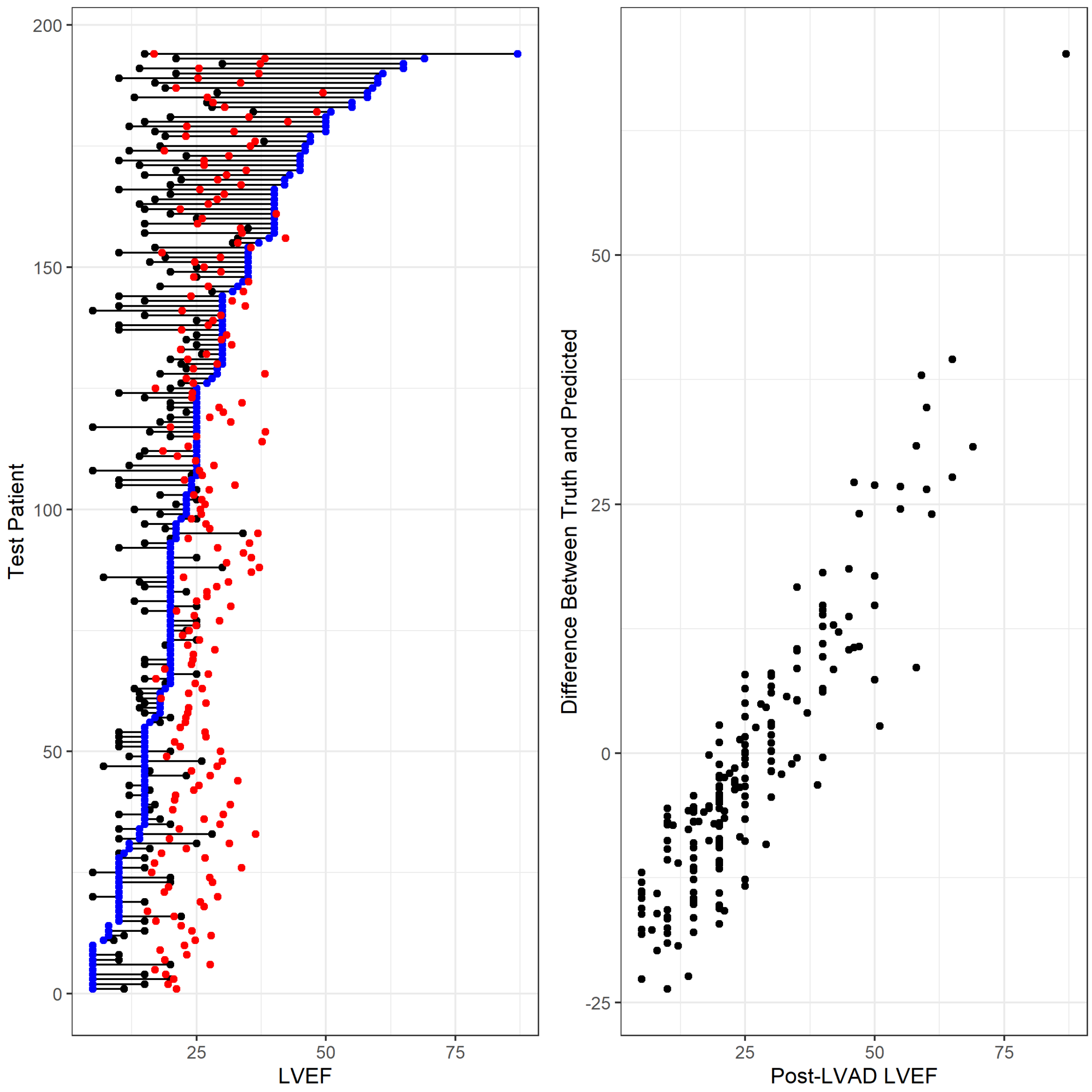


**Supplemental Figure 3**: Visual Representation of baseline-, post-, and predicted-, left ventricular ejection fraction (LVEF) (left). Plots shown are of baseline (black circles), true post-LVAD: LVEF (blue circles), and predicted post-LVAD: LVEF (red circles) for each of the testing dataset patients (n=69). Predictions are conservative for the patients with the most improvement. Bland-Altman Plots (right) show the difference between true LVEF (patients post-LVAD: LVEF) and predicted value. The solid line represents the average difference between truth and predicted, while the dashed lines represent 95% confidence intervals. Model performance diminishes as the average of the truth and predicted move to the extremes (either minimal or exaggerated change in LVEF).

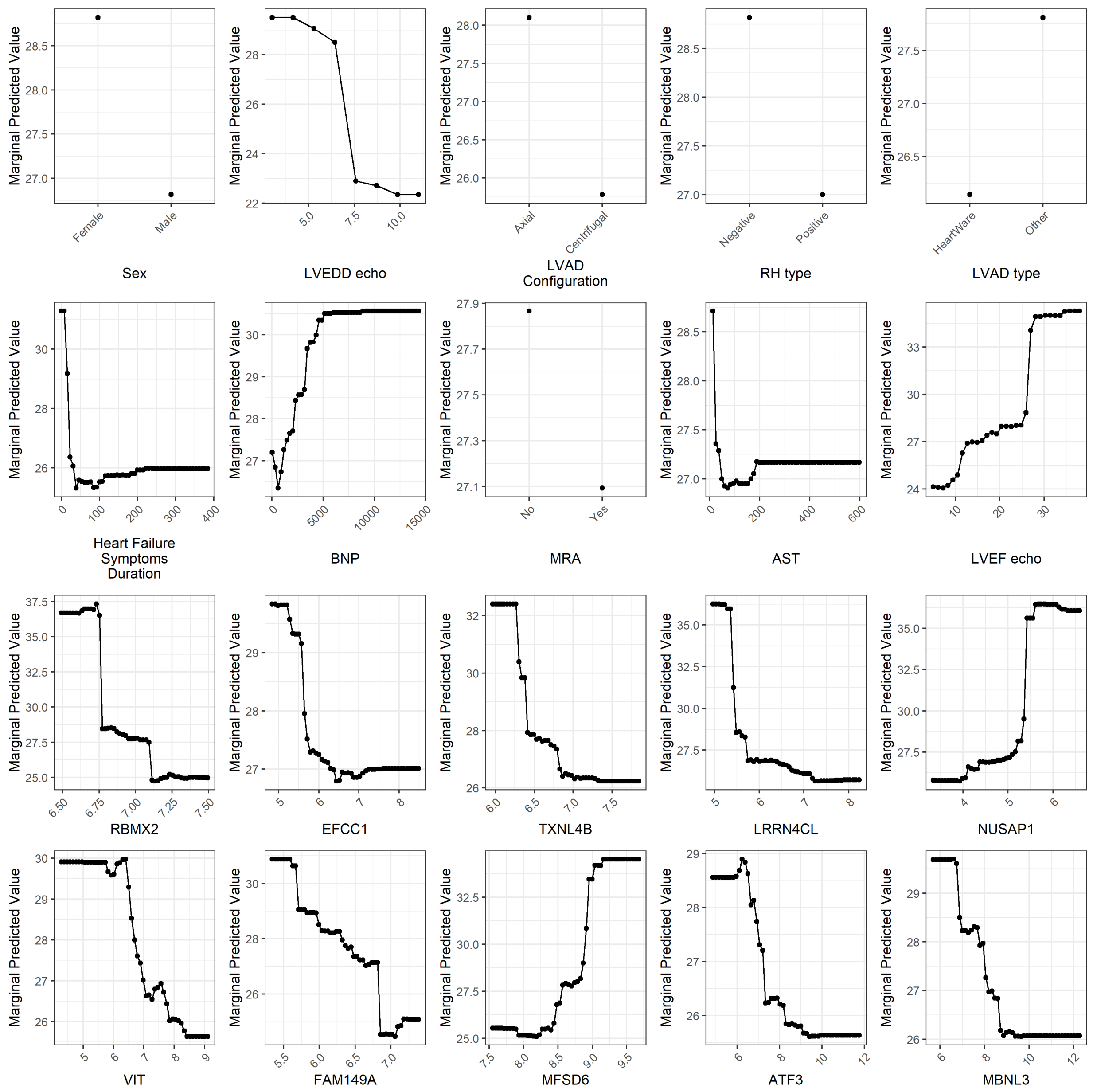


**Supplemental Figure 4**: Partial Dependency Plots for the prediction of improved myocardial function (LVEF). From the Random Forest (rf) two-stage screening process, the top 10 clinical and mRNA-transcript variables are shown. Y-Axis: An increased marginal predicted value for LVEF indicates a greater probability for a variable to predict LVAD-mediated functional myocardial improvement. X-Axis: Sex (Female vs. Male), LVEDD echo (cm), LVAD Configuration (Axial vs. Centrifugal), RH type (Negative vs. Positive), LVAD type (HeartWare vs. Other), Heart Failure Symptoms Duration (months), BNP (pg/mL), MRA (No vs. Yes), AST (U/L), LVEF echo (%), *RBMX2*, *EFCC1*, *TXNL4B*, *LRRN4CL*, *NUSAP1*, *VIT*, *FAM149A*, *MFSD6*, *ATF3*, *MBNL3* (gene expression represented by R-Log Values).

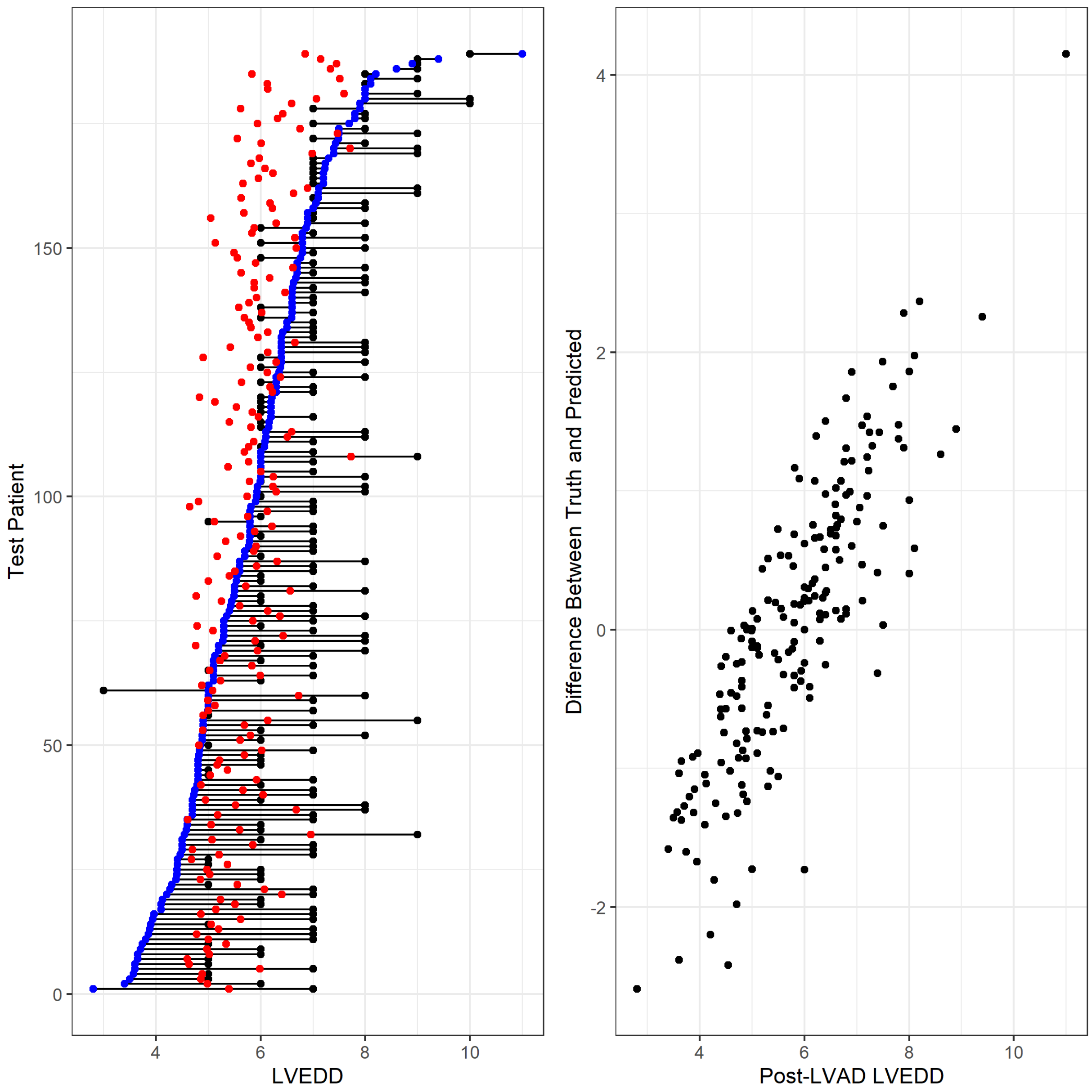


**Supplemental Figure 5:** Visual Representation of baseline-, post-, and predicted-, left ventricular end-diastolic diameter (LVEDD) (left). Plots shown are of baseline (black circles), true post-LVAD: LVEDD (blue circles), and predicted post-LVAD: LVEDD (red circles) for each of the testing dataset patients (n=69). The model was conservative in predicting improved LVEDD, underestimated the decrease in LVEDD when the true change from baseline was great. Bland-Altman Plots (right) show the difference between true LVEDD (patients post-LVAD: LVEDD) and predicted value. The solid line represents the average difference between truth and predicted, while the dashed lines represent 95% confidence intervals. Model performance diminishes as the average of the truth and predicted move to the extremes (either minimal or exaggerated change in LVEDD).

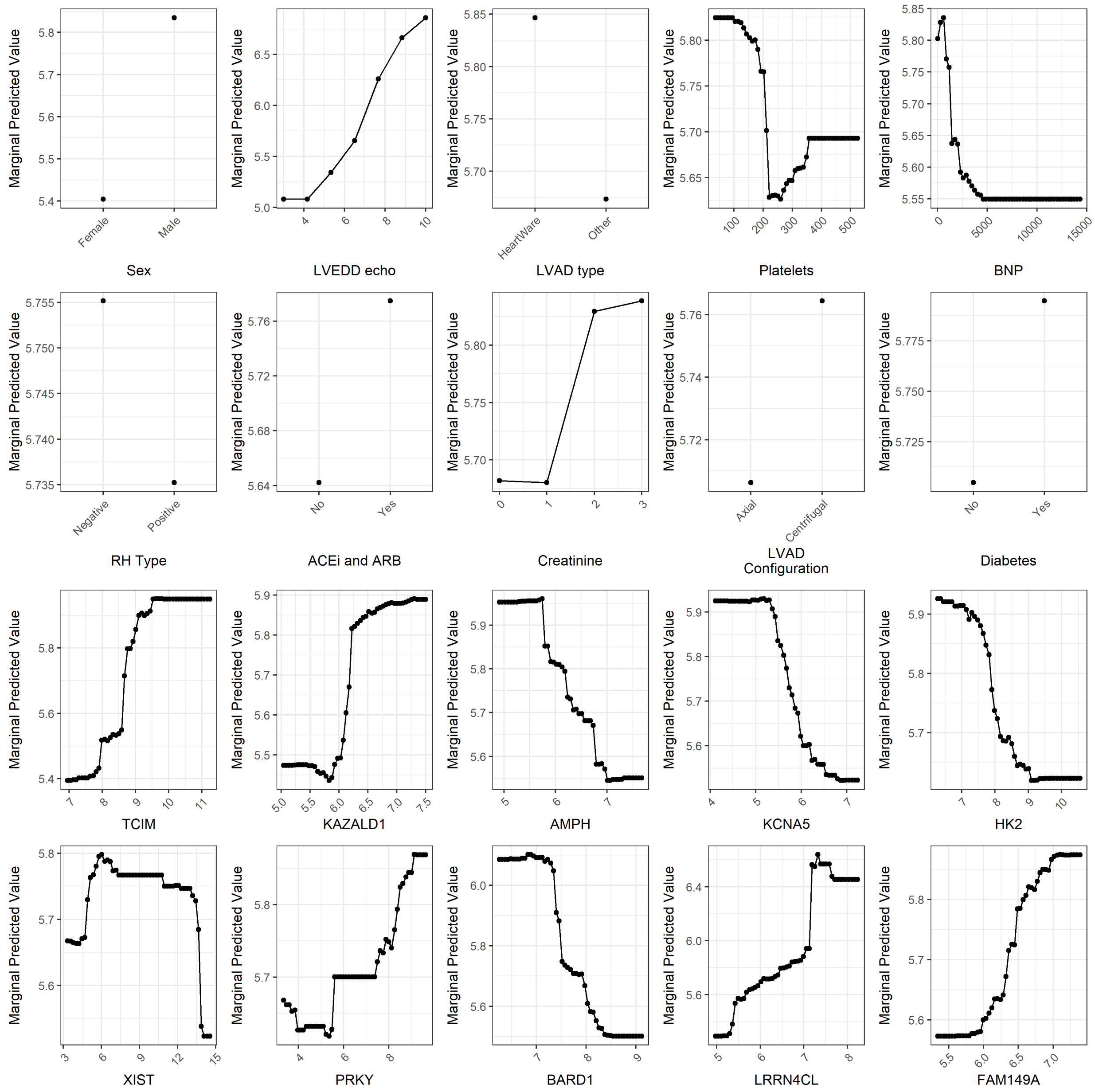


**Supplemental Figure 6**: Partial Dependency Plots for the prediction of improved myocardial structure (LVEDD). From the Random Forest (rf) two-stage screening process, the top 10 clinical and mRNA-transcript variables are shown. Y-Axis: A decreased marginal predicted value for LVEDD indicates a greater probability for a variable to predict LVAD-mediated structural myocardial improvement. X-Axis: Sex (Female vs. Male), LVEDD echo (cm), LVAD type (HeartWare vs. Other), Platelets (mcL), BNP (pg/mL), RH type (Negative vs. Positive), ACEi and ARB (No vs. Yes), Creatinine (mg/dL), LVAD Configuration (Axial vs. Centrifugal), Diabetes (No vs. Yes), *TCIM*, *KAZALD1*, *AMPH*, *KCNA5*, *HK2*, *XIST*, *PRKY*, *BARD1*, *LRRN4CL*, *FAM149A* (gene expression represented by R-Log Values).

**Supplemental Table 1:** Clinical characteristics of patients by institution.

| Variable | All patients (N=208) | Allegheny (N=53) | Louisville (N=32) | UCAR (N=123) | P-value |
|---|---|---|---|---|---|
| ***Demographics*** | | | | | |
| Age, years | 55.7 (15.2) | 58.7 (12.1) | 50.9 (14.2) | 55.7 (16.3) | 0.07[a] |
| Male sex, n (%) | 164 (79%) | 40 (75.5%) | 24 (75%) | 100 (81.3%) | 0.58[c] |
| Race | | | | | |
| White, n (%) | 179 (86.1%) | 46 (86.8%) | 27 (84.4%) | 106 (86.2%) | 0.67[f] |
| Black, n (%) | 15 (7.2%) | 4 (7.5%) | 5 (15.6%) | 6 (4.9%) | 0.10[f] |
| Other, n (%) | 14 (6.7%) | 3 (5.7%) | 0 (0%) | 11 (8.9%) | 1.00[f] |
| Body Mass Index, kg/m$^2$ | 28.8 (5.8) | 29.1 (6.1) | 27.9 (4.8) | 28.9 (5.9) | 0.66[a] |
| Blood Type | | | 0.64[f] | | 1.00[f] |
| A, n (%) | 46 (37%) | 1 (50%) | 0 (NaN%) | 45 (36.6%) | |
| B, n (%) | 20 (16%) | 0 (0%) | 0 (NaN%) | 20 (16.3%) | |
| AB, n (%) | 4 (3%) | 0 (0%) | 0 (NaN%) | 4 (3.3%) | |
| O, n (%) | 55 (44%) | 1 (50%) | 0 (NaN%) | 54 (43.9%) | |
| Rhesus D Antigen | | | | | 0.52[f] |
| Negative, n (%) | 27 (15%) | 6 (19.4%) | 3 (9.4%) | 18 (14.6%) | |
| ***Medical History*** | | | | | |
| Smoking, n (%) | 109 (53%) | 38 (74.5%) | 17 (53.1%) | 54 (43.9%) | 0.001[c] |
| Diabetes Mellitus, n (%) | 69 (33%) | 18 (34.6%) | 11 (34.4%) | 40 (32.8%) | 0.97[c] |
| Hypertension, n (%) | 112 (55%) | 33 (66%) | 24 (75%) | 55 (44.7%) | 0.002[c] |
| Ischemic Cardiomyopathy, n (%) | 71 (36%) | 19 (43.2%) | 14 (43.8%) | 38 (30.9%) | 0.20[c] |
| New York Heart Association Classification | | | | | 0.008[c] |
| 3, n (%) | 60 (29%) | 8(15.4%) | 15(46.9%) | 37(30.1%) | |
| 4, n (%) | 147 (71%) | 44(84.6%) | 17(53.1%) | 86(69.9%) | |
| Heart Failure Symptoms Duration, months | 71.1 (71.6) | 64.9 (65.9) | 30.1 (39.5) | 84.5 (76.3) | <0.001[a] |
| Cardiac Resynchronization/Implantable Cardioverter Defibrillator, n (%) | 172 (83%) | 38 (74.5%) | 27 (84.4%) | 107 (87%) | 0.05[c] |
| INTERMACS profile | | | | | <0.001[a] |
| 1, n (%) | 27 (16%) | 16 (32%) | 0 (NaN%) | 11 (8.9%) | |
| 2, n (%) | 36 (21%) | 16 (32%) | 0 (NaN%) | 20 (16.3%) | |
| 3, n (%) | 58 (34%) | 10 (20%) | 0 (NaN%) | 48 (39%) | |
| 4-7, n (%) | 52 (31%) | 8 (16%) | 0 (NaN%) | 44 (35.8%) | |
| Left Ventricular Assist Device Type | | | | | <0.001[s] |
| HeartMate II, n (%) | 123 (60%) | 32 (64%) | 23(71.9%) | 68 (55.3%) | |
| HeartMate 3, n (%) | 13 (6%) | 10 (20%) | 0 (0%) | 3 (2.4%) | |
| HeartWare, n (%) | 55 (27%) | 8 (16%) | 5(15.6%) | 42 (34.1%) | |
| Other, n (%) | 14 (6%) | 0 (0%) | 4 (12.4%) | 10 (8.1%) | |
| Left Ventricular Assist Device Configuration | | | | | 0.10[c] |
| Axial, n (%) | 133 (66%) | 32 (64%) | 25 (83.3%) | 76 (62.8%) | |
| Left Ventricular Assist Device Support Duration, days | 565.3 (640.8) | 421.0 (405.4) | 0 (NaN%) | 571.0 (649.1) | 0.65[a] |
| ***Pre-operative Supportive Therapies*** | | | | | |
| Inotrope Dependency, n (%) | 136 (68%) | 29 (64.4%) | 28 (87.5%) | 79 (64.2%) | 0.036[c] |
| Intra-aortic Balloon Pump, n (%) | 30 (15%) | 18 (39.1%) | 5 (15.6%) | 7 (5.7%) | <0.001[f] |
| Temporary Mechanical Circulatory Support, n (%) | 8 (4%) | 3 (6.7%) | 1 (3.1%) | 4 (3.3%) | 0.52[f] |
| ***Heart Failure Medical Therapy*** | | | | | |
| Beta-Blockers, n (%) | 142 (70%) | 25(52.1%) | 23 (71.9%) | 94 (76.4%) | 0.007[c] |
| Angiotensin Converting Enzyme Inhibitors/Angiotensin Receptor Blockers, n (%) | 85 (42%) | 23(46.9%) | 9(28.1%) | 53(43.1%) | 0.09[a] |
| Mineralocorticoid Receptor Antagonists, n (%) | 106 (63%) | 26 (57.8%) | 0 (NaN%) | 80 (65%) | 0.47[f] |

| | | | | | |
|---|---|---|---|---|---|
| Diuretics, n (%) | 151 (90%) | 34 (77.3%) | 0 (NaN%) | 117 (95.1%) | 0.001[f] |
| ***Right Heart Catheterization*** | | | | | |
| Right Atrial Pressure, mmHg | 11.9 (6.3) | 12.5 (7.0) | 10.4 (5.9) | 12.0 (6.2) | 0.35[a] |
| Pulmonary Capillary Wedge Pressure, mmHg | 23.8 (8.3) | 23.2 (8.3) | 21.6 (7.1) | 24.5 (8.5) | 0.23[a] |
| Cardiac Index, L/min/$m^2$ | 1.9 (0.6) | 1.8 (0.6) | 2.2 (0.6) | 1.9 (0.6) | 0.015[a] |
| ***Laboratory Values*** | | | | | |
| White Blood Cells, $x10^9$/µL | 9.1 (3.6) | 10.3 (3.9) | 12.3 (2.3) | 8.1 (3.4) | <0.001[a] |
| Platelets, $x10^9$/µL | 209.1 (69.7) | 216.5 (59.9) | 210.5 (60.7) | 208.3 (72.7) | 0.95[a] |
| Hemoglobin, g/dl | 12.5 (2.2) | 9.8 (1.3) | 0 (NaN%) | 12.6 (2.2) | 0.002[a] |
| Sodium, mEq/L | 134.7 (4.8) | 135.2 (5.2) | 135.4 (3.8) | 134.4 (5.1) | 0.61[a] |
| Blood Urea Nitrogen, mg/dL | 26.9 (13.9) | 19.2 (8.4) | 25.9 (14.6) | 27.6 (13.9) | 0.32[a] |
| Creatinine, mg/dL | 1.3 (0.6) | 0.8 (0.4) | 1.4 (0.6) | 1.3 (0.5) | 0.06[a] |
| Total Bilirubin, mg/dL | 1.6 (1.2) | 1.2 (0.4) | 0 (NaN%) | 1.6 (1.2) | 0.39[a] |
| Albumin, g/dL | 3.7 (0.6) | 3.5 (0.5) | 3.5 (0.6) | 3.8 (0.6) | 0.022[a] |
| B-type Natriuretic Peptide, pg/mL | 1469.1 (1938.3) | 1329.4 (2137.0) | 2039.7 (3240.9) | 1362.1 (1182.8) | 0.19[a] |
| ***Baseline Echocardiographic Parameters*** | | | | | |
| Left Ventricular Ejection Fraction, % | 17.6 (6.8) | 14.1 (6.4) | 16.9 (6.4) | 19.2 (6.5) | <0.001[a] |
| Left Ventricular End-Diastolic Diameter, cm | 6.9 (1.2) | 6.9 (1.3) | 7.6 (0.9) | 6.7 (1.1) | <0.001[a] |
| ***Post-LVAD Echocardiographic Parameters*** | | | | | |
| Left Ventricular Ejection Fraction, % | 27.3 (15.4) | 21.1 (15.5) | 32.7 (20.5) | 28.6 (13.0) | <0.001[a] |
| Left Ventricular End-Diastolic Diameter, cm | 5.7 (1.3) | 5.8 (1.4) | 6.2 (1.7) | 5.6 (1.1) | 0.09[a] |

[a] ANOVA, [c] Chi-squared test, [f]Fisher's exact test, [s] Chi-squared test by Montecarlo simulation.
